# Benchmarking cell type annotation in spatial transcriptomics: resolving cellular hierarchies, biological fidelity, and dynamic cell states

**DOI:** 10.64898/2026.06.16.732716

**Authors:** Yuling Zhu, Yunfei Hu, Manfei Bella Xie, Haoran Qin, Zuzanna J Szul, Derek M Young, Weiman Yuan, Qimeng Wang, Yichen Henry Liu, Wenjun Shen, Shan Meltzer, Xin Maizie Zhou

## Abstract

Spatial transcriptomics enables the quantification of gene expression within its native tissue context, providing unprecedented insight into tissue architecture, cellular ecosystems, and local cell–cell interactions at regional and single-cell resolution. Accurate cell type annotation is a critical prerequisite for interpreting these data and is often the first and most essential step in downstream analysis. Despite rapid advances in computational methods, cell type annotation remains challenging and frequently requires extensive expert-driven manual curation based on marker-gene expression, spatial context, and prior biological knowledge. While early approaches relied primarily on transcriptional similarity, newer methods increasingly incorporate spatial information, histological features, and multimodal data to improve annotation accuracy. Nevertheless, reliable annotation remains difficult when biological interpretation requires fine-grained subtype resolution, particularly for platforms with limited gene panels, tissues undergoing dynamic cellular state transitions, and studies in which reference and query datasets differ substantially in biological context or technical modality. Here, we present a systematic benchmark of 20 state-of-the-art cell type annotation methods across four spatial transcriptomics datasets spanning diverse technologies, experimental conditions, cell numbers, and gene panel sizes. Importantly, all benchmark datasets contain expert-curated cell type labels, including well-resolved cell populations and subtype annotations, providing high-quality biological ground truth for evaluation. The benchmark encompasses both reference-based and reference-free methods representing a broad range of computational frameworks. Performance was assessed using conventional classification metrics, including accuracy and F1-based measures, together with structure-aware metrics that evaluate both cell-level annotation accuracy and preservation of higher-order biological organization. Across datasets, annotation performance varied substantially according to tissue context, reference–query similarity, and annotation granularity. Fine-grained subtype annotation and recovery of rare cell populations remained challenging for many methods, particularly in datasets capturing injury, repair, developmental, and regenerative processes characterized by continuous cellular state transitions. Notably, high classification accuracy did not necessarily correspond to preservation of global cellular relationships or biologically coherent downstream pathway and gene-set enrichment analyses. Overall, scANVI, Seurat, and TACCO consistently ranked among the top-performing methods, although their relative advantages were context dependent. Together, our results provide a comprehensive assessment of current annotation strategies for spatial transcriptomics and offer practical guidance for selecting methods that best align with specific biological questions, dataset characteristics, and analytical priorities.

## Introduction

Advances in spatial transcriptomics (ST) technologies enable transcriptome-wide molecular profiling *in situ* while preserving native tissue architecture and spatial cellular organization [1, 2]. By integrating gene expression with spatial context, ST provides a powerful framework for studying tissue architecture, cellular interactions, and microenvironmental organization across development, physiology, and disease. These technologies have substantially expanded our ability to characterize how distinct cell populations are spatially arranged [3, 4, 5], interact with neighboring cells, and collectively contribute to tissue function and pathology.

A central step in nearly all spatial transcriptomic analyses is accurate cell type annotation. Reliable identification and localization of cell types form the basis for downstream analyses, including spatial domain identification, tissue niche characterization, cellcell communication inference, and disease-associated microenvironment discovery [6]. In particular, highresolution annotation at the subtype or cellular-state level is essential for resolving fine-grained tissue organization and detecting rare or spatially restricted populations that may play disproportionate functional roles. As spatial transcriptomics technologies continue to improve in throughput and resolution, the demand for accurate and scalable cell type annotation methods has become increasingly important.

Despite rapid methodological advances, accurate subtype-level annotation in ST data remains challenging. Closely related cell states frequently exhibit subtle transcriptional differences that are difficult to distinguish in spatial measurements. These challenges are further compounded by technical variability across platforms, sparse transcript capture, and mixed-cell-resolution measurements in which expression signals from multiple neighboring cells can overlap [1]. In addition, heterogeneous tissue architectures and spatially varying cellular states can reduce the robustness of existing label transfer approaches, particularly when reference single-cell datasets incompletely capture the diversity of spatial cell states. Together, these limitations hinder precise characterization of cellular heterogeneity and spatial organization in complex tissues.

Based on whether reference datasets are required, existing cell type annotation methods can be broadly categorized into reference-based and reference-free approaches. From a methodological perspective, these methods span a wide range of mathematical frameworks, including regression-based models [7, 8], dimensionality reduction techniques [9], anchor-based nearest neighbor frameworks [10], optimal transport [11], correlation-based approaches [12], optimization-based frameworks [13] and deep generative models [14].

In recent years, increasingly complex models have emerged, including large-scale transcriptome foundation models such as scGPT [15] and Nicheformer [16], as well as large language model–based approaches such as GPTCelltype [17]. Unlike traditional annotation methods that rely primarily on explicit reference matching or feature engineering, transcriptome foundation models are pretrained on large collections of single-cell or spatial transcriptomic datasets to learn biologically meaningful and transferable representations. The resulting representations provide a robust basis for downstream annotation tasks via head tuning or task-specific fine-tuning on labeled data [16]. In parallel, LLM-based approaches such as GPTCelltype utilize marker gene information to generate freetext cell type labels, leveraging the extensive biological knowledge and reasoning capabilities encoded in large pretrained language models to support flexible cell type annotation across diverse tissues and experimental settings [17].

More recently, Crowley et al. introduced AnnDictionary, a unified framework that leverages large language models for de novo cell type and gene set annotation, and systematically benchmarked a wide range of commercial LLMs on the Tabula Sapiens atlas [18]. Their results demonstrated that LLMs can recover broad cell type identities with high semantic agreement with manual annotations, particularly for abundant and well-characterized cell populations. However, these studies primarily focus on open-vocabulary cell type annotation through natural language label generation and are not designed to systematically assess annotation accuracy under varying reference quality, cell type resolution mismatch, or cross-dataset generalization factors that remain central challenges in the benchmarking and evaluation of cell type annotation methods.

Furthermore, a systematic assessment of how well current computational methods resolve fine-grained cell type identities in spatial transcriptomics datasets remains lacking. For sequencing-based spatial transcriptomics platforms that have not yet achieved single-cell resolution, several comprehensive benchmarking studies [6, 19, 20] have evaluated methods for inferring cell type composition at individual spatial locations. These efforts primarily focus on spot-level deconvolution accuracy and assess performance using aggregate metrics such as overall accuracy or correlation-based measures. However, considerably less attention has been devoted to assessing annotation performance across different levels of the cell type hierarchy, particularly among closely related cell subtypes. As a result, it remains unclear how effectively existing methods can resolve subtle cellular heterogeneity in spatial contexts. A related limitation is that most benchmarking studies rely on aggregate performance metrics, such as overall accuracy, without accounting for the hierarchical relationships among cell types [21]. Consequently, all misclassifications are treated equally, regardless of whether a method confuses two distinct major cell classes or merges closely related subtypes within the same lineage. In practice, some methods may fail to distinguish subtle subpopulations while still correctly assigning cells to their broader lineage, a biologically meaningful outcome that conventional accuracy metrics do not adequately capture.

Although many widely used cell type annotation methods were originally developed for single-cell transcriptomics data and rely primarily on each cell’s molecular profile, an increasing number of recent approaches have begun to incorporate spatial context into the annotation process. Nevertheless, a cell’s identity and functional state are often influenced by its local microenvironment, including interactions with neighboring cells, tissue architecture, and spatially organized signaling cues [22]. As spatial transcriptomics technologies continue to mature, accumulating evidence suggests that incorporating neighborhood information can improve the identification of cell states and resolve cellular heterogeneity that may be difficult to distinguish from transcriptomic profiles alone. At the same time, the limited availability of high-quality annotated spatial transcriptomics datasets and the substantial domain shifts across tissues, technologies, and experimental protocols remain major obstacles to robust model generalization. Recent studies have shown that deep learning–based cell annotation models can experience significant performance degradation when applied to datasets that differ from their training distributions [23]. Moreover, many existing cell representation learning approaches are still primarily optimized for gene-expression similarity and may not fully capture spatial dependencies, cell–cell interactions, or tissue-level organization.

To address these limitations, a growing body of work has introduced spatially aware annotation frameworks that leverage neighborhood aggregation, graph neural networks, spatially informed representation learning, and foundation-model architectures to incorporate cellular context into learned embeddings. Despite these methodological advances, it remains unclear under which conditions explicit spatial modeling provides measurable improvements in annotation performance, how robust these gains are across datasets and technologies, and whether spatial context is particularly beneficial for resolving finegrained cell subtypes or ambiguous cellular states.

Consequently, there is an urgent need for a systematic evaluation that addresses these outstanding challenges and provides a comprehensive assessment of current cell type annotation methods. In particular, the performance of existing annotation tools across spatial transcriptomics technologies, reference–query mismatches, hierarchical levels of cell identity, and biologically dynamic conditions has not been systematically characterized. Existing benchmarks have primarily focused on spot-level deconvolution or aggregate annotation accuracy, leaving the evaluation of fine-grained subtype annotation largely unexplored. In particular, few studies distinguish between broad lineage misclassification and confusion among closely related subtypes, or systematically assess whether explicit spatial modeling improves annotation accuracy in real-world spatial transcriptomics datasets. Furthermore, the robustness of annotation methods under cross-platform and cross-condition transfer remains poorly understood, especially when reference and query data differ in tissue state, developmental stage, disease condition, or experimental technology. Although LLM-based approaches have demonstrated promise in translating marker genes signatures into interpretable cell type labels, they complement, rather than replace quantitative annotation frameworks and do not directly address these benchmarking challenges.

Here, we present a comprehensive benchmark of a broad collection of cell type annotation methods across four spatial transcriptomics datasets spanning multiple technologies and biological contexts. Importantly, all four datasets contain expert-curated cell type annotations, providing a high-quality biological ground truth for evaluation. We evaluated reference-based, reference-free, spatially aware, foundationmodel-based, and LLM-assisted approaches using a diverse set of performance measures, including classification, clustering, biological structure preservation, hierarchical annotation, and marker-enrichment metrics. Our benchmark reveals substantial variation in method performance across datasets, cell type resolutions, and reference-query settings. We show that fine-grained cell subtype annotation remains considerably more challenging than broad lineage assignment, even when higher-level cell identities are accurately recovered. Furthermore, the benefits of incorporating spatial information are context dependent rather than universal, while foundation model and LLM-based approaches exhibit promising performance but do not consistently outperform established annotation methods across all evaluation scenarios. Together, these results provide a systematic framework for evaluating annotation accuracy across multiple biological and technical dimensions and offer practical guidance for method selection and interpretation in spatial transcriptomics studies.

## Results

### The overall benchmarking framework

In this study, we established a systematic benchmarking framework to evaluate cell type annotation methods for spatial transcriptomics across diverse experimental platforms and analytical settings (Fig. 1). Our pipeline begins with four spatial transcriptomics datasets [2, 24, 25, 26], together with matched or external single-cell/single-nucleus RNA-seq references when required. We then evaluated a broad collection of cell type annotation methods spanning diverse methodological paradigms (Fig. 1a and Table 1). These included 17 reference-based approaches, encompassing foundation models (scGPT [15], scCello [27], Nicheformer [16], and SToFM [28]), deep learning models (scANVI [14], Spatial-ID [29], and GraphST [30]), probabilistic frameworks (DestVI [31], Cell2location [1], and RCTD [32]), dimensinality-reduction-based methods (CARD [9]), optimal transport approaches (TACCO [11]), regression-based methods (SpatialDWLS [7] and SPOTlight [8]), anchor-based nearest-neighbor methods (Seurat [10]), correlation-based approaches (SingleR [12]), and optimization-based frameworks (Tangram [13]). We additionally evaluated three reference-free methods STdeconvolve [33], SpaGCN [3], and BANKSY [34] to assess annotation performance in the absence of external reference data. Parameter settings for all benchmarked tools are listed in Additional file 1: Supplementary Table 1. Brief descriptions of each method are provided in the Methods section. The supervised training strategies used for foundation models, including reference mapping, head tuning, and partial fine tuning are illustrated alongside the corresponding models.

**Table 1:**
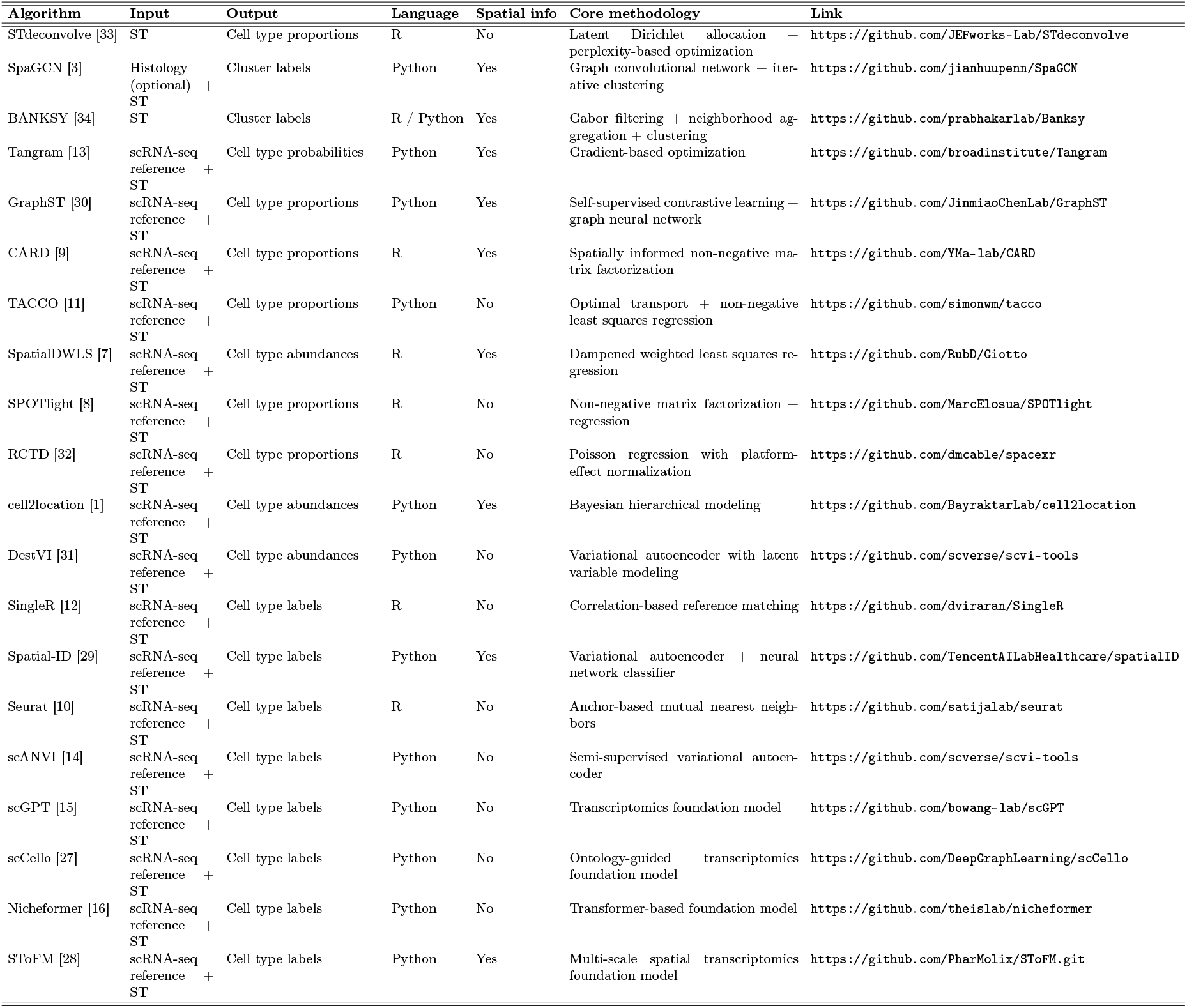
Overview of the 20 cell type annotation methods evaluated in this study, categorized by input requirements, output type, programming language, utilization of spatial information, and core methodological approach.

**Figure 1:**
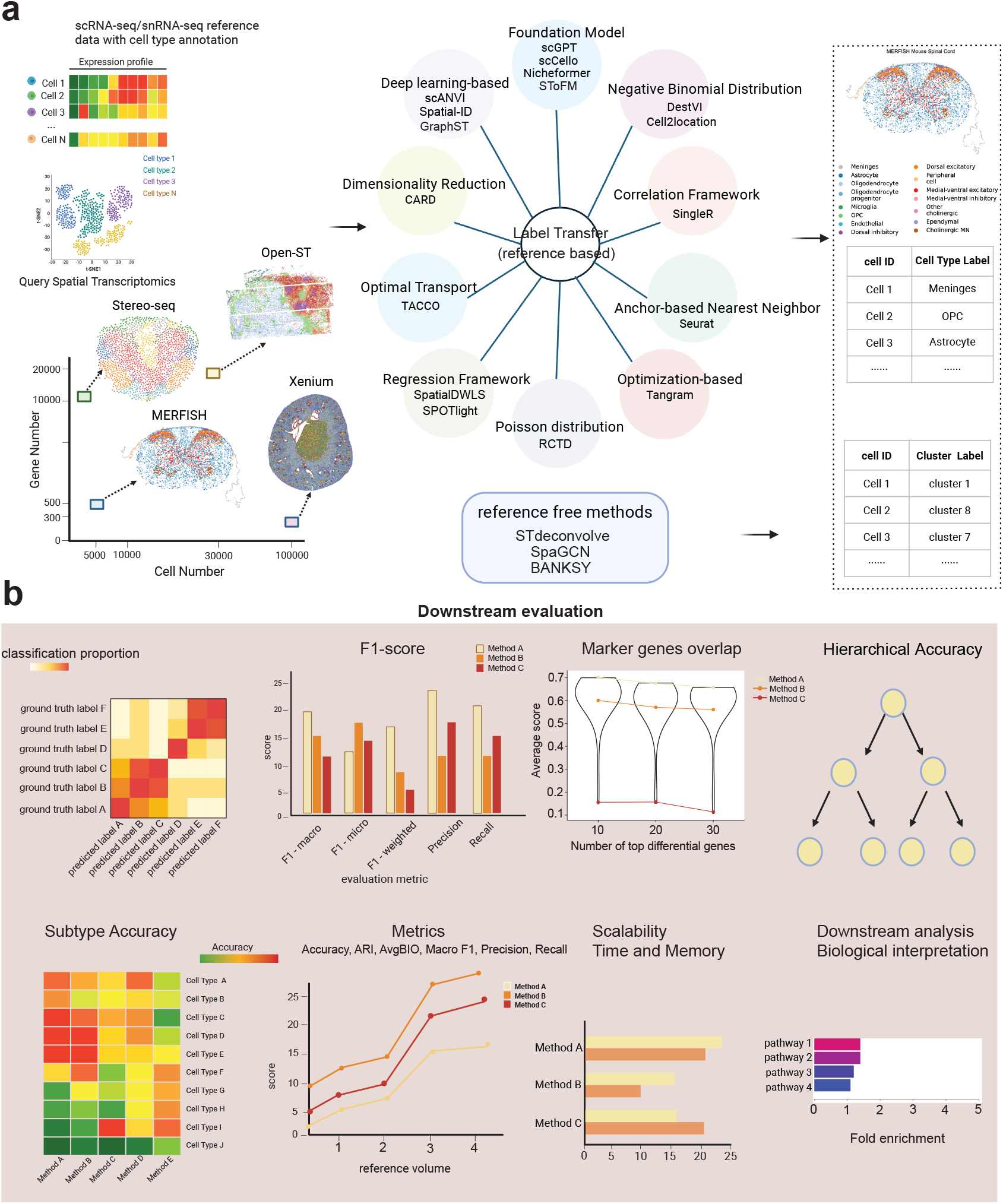
Overview of the benchmarking pipeline for cell type annotation methods and downstream evaluation on spatial transcriptomics data. **(a)** Benchmarking workflow. Spatial transcriptomics datasets generated by multiple platforms, including MERFISH, Open-ST, Stereo-seq, and Xenium, were analyzed using both reference-based label transfer methods and reference-free methods. For referencebased annotation, annotated single-cell RNA-seq/single-nucleus RNA-seq datasets were used as references to transfer cell type labels to query spatial transcriptomics cells. **(b)** Predicted labels were assessed against ground-truth annotations using multiple complementary criteria. Classification performance was evaluated using confusion matrix, quantitative metrics, hierarchical accuracy, subtype-level accuracy, scalability in runtime and memory, robustness to reference volume, and biological concordance through downstream analyses such as marker-gene overlap and pathway enrichment. Together, this framework provides a systematic assessment of both annotation accuracy and biological interpretability across methods.

Among the 20 benchmarked methods, nine methods explicitly or implicitly incorporate spatial information, although they differ markedly in how spatial context is modeled. SpatialDWLS uses spatially informed preprocessing based on k-nearestneighbor clustering [7], whereas SpaGCN constructs a graph integrating gene expression, spatial coordinates, and histology, followed by graph convolution to aggregate information from neighboring cells [3]. BANKSY augments each cell’s expression profile with a spatial-kernel-weighted representation of its neighbors and performs clustering in this expanded product space [34]. Other methods, such as GraphST and CARD, incorporate spatial information through graph-based representations, spatial regularization, or neighborhood-aware embedding learning.

The resulting predicted cell labels or clusters were compared with curated ground truth annotations using a multi-dimensional evaluation framework, including global classification performance, subtype-level accuracy, hierarchical annotation accuracy, marker-gene concordance, computational scalability, and downstream biological interpretability (Fig. 1b). This design enabled us to evaluate not only overall annotation accuracy but also method robustness, transcriptional fidelity, sensitivity to reference selection, and the biological validity of downstream analyses.

To instantiate this benchmarking framework, we selected four high-quality spatial transcriptomics datasets (Table 2: MERFISH mouse spinal cord dataset [24], Open-ST human lymph node dataset [25], Xenium kidney dataset [2], and Stereo-seq axolotl brain dataset [26]) with expert-curated cell type annotations serving as biological ground truth. These datasets were generated using diverse technologies and span a wide range of gene coverage, cell numbers, and spatial resolutions. The diversity of these datasets captures substantial variation in transcriptome capture sensitivity, spatial resolution, and biological complexity across current spatial transcriptomics platforms. To comprehensively evaluate method performance under biologically challenging and realistic scenarios, we designed six reference-query experimental settings. For the MERFISH mouse spinal cord dataset, 17 tissue sections from the same study were used as a matched reference. In a second MERFISH setting, focused on mouse spinal cord data, the same test section was annotated using an external single-nucleus RNA-seq reference. Similarly, the Xenium mouse kidney dataset, comprising 12 sections, was evaluated with an external single-nucleus RNA-seq reference. To assess crosssection annotation performance, the Open-ST human lymph node dataset used one tissue section for training and a serial section from the same individual for testing. Finally, to evaluate performance in dynamic biological systems characterized by substantial spatial reorganization, we analyzed the Stereo-seq axolotl brain dataset under two conditions. In the regeneration setting, three sections collected at 5 days post-injury were used, with two sections serving as the reference and the third as the query. In the developmental setting, a later developmental stage (stage 54) was used as the reference to annotate an earlier stage (stage 44). Together, these datasets and experimental settings capture a broad range of biological and technical heterogeneity commonly encountered in real-world spatial transcriptomics studies.

**Table 2:** Characteristics of the spatial transcriptomics datasets used for benchmarking, including ST type, experimental protocol, sample size (cells and genes), number of tissue sections, and data source.

| Dataset | ST type | Protocol | Cells / Genes | Num. of slices | Source |
| --- | --- | --- | --- | --- | --- |
| Mouse spinal cord | In situ capture | MERFISH | 41267 / 500 | 18 | [https://www.cell.com/cell/fulltext/S0092-8674(24)00578-1](https://www.cell.com/cell/fulltext/S0092-8674%2824%2900578-1) |
| Human metastatic lymph node | In situ capture | Open-ST | 77722 / 28943 | 2 | https://support.10xgenomics.com/spatial-gene-expression/datasets |
| Mouse six stages renal injury and repair | In situ capture | Xenium | 1374915 / 300 | 12 | [https://www.cell.com/cell/fulltext/S0092-8674(24)00578-1](https://www.cell.com/cell/fulltext/S0092-8674%2824%2900578-1) |
| Axolotl brain development and regeneration | In situ capture | Stereo-seq | 4406 / 12704<br>24656 / 27184 | 5 | https://db.cngb.org/stomics/artista/download/ |

### Comparative performance of cell type annotation methods across datasets

Previous benchmark studies have demonstrated that no single domain identification tool consistently achieves the best performance across diverse datasets and analytical scenarios [5]. Instead, performance is often highly context-dependent, varying according to data modality, tissue type, biological complexity, and experimental design. Building on these observations, we evaluated each method across three complementary dimensions classification accuracy, preservation of tissue organization and biological coherence, and computational scalability to provide scenariospecific guidance for users with diverse analytical objectives.

We employed a comprehensive set of metrics spanning three evaluation dimensions (Fig. 2). For classification accuracy, we calculated Accuracy alongside Macro, Micro, and Weighted F1 scores. Because Micro F1 is mathematically equivalent to overall Accuracy in our single-label multiclass setting, we use Accuracy as the primary performance metric throughout. To assess preservation of tissue organization and biological coherence, we evaluated tissue architecture using Element-Centric Similarity (ECS), Adjusted Rand Index (ARI), Normalized and Adjusted Mutual Information (NMI and AMI), Average Silhouette Width (ASW), and the Average Biological Conservation score (AvgBIO). Details for these metrics are provided in the Methods section. For computational scalability, we measured execution time and peak memory usage.

**Figure 2:**
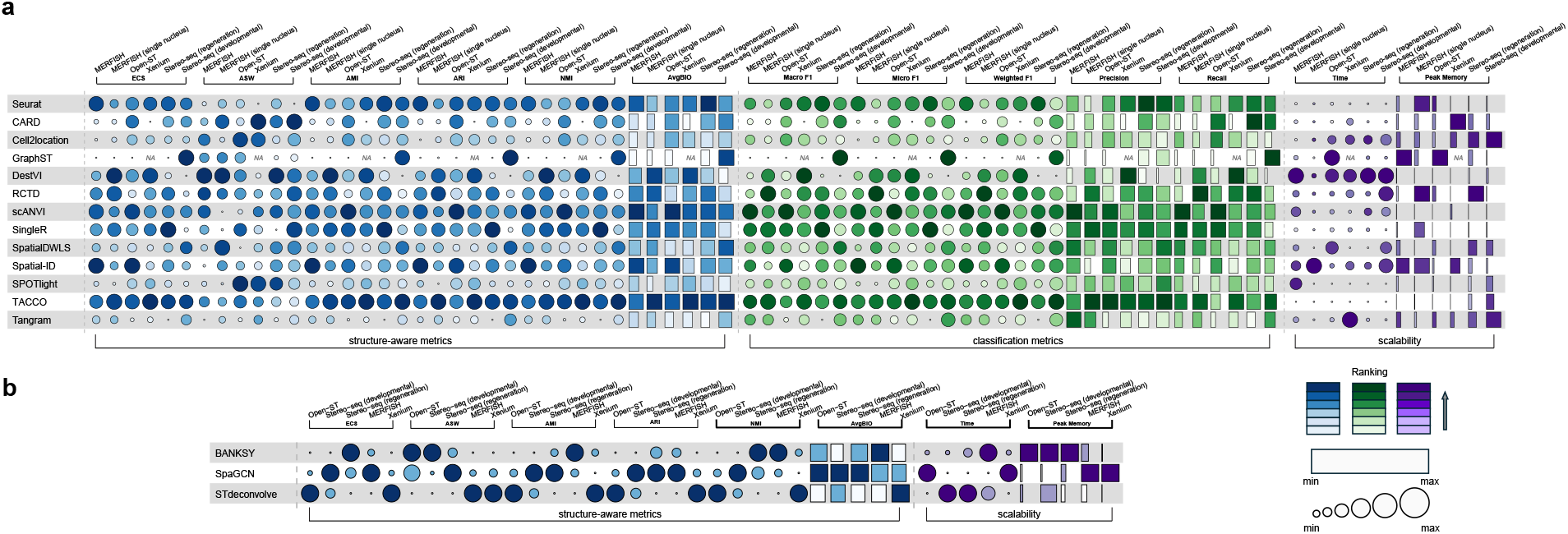
Summary of method performance across all datasets. Classification metrics are shown in green, structure-aware metrics in blue, and scalability metrics in purple. Darker shades indicate better performance for classification and structure-aware metrics, but greater computational cost for scalability metrics. Larger circles and longer rectangles indicate larger metric values relative to the other methods within the same metric–dataset column. Symbol sizes were scaled separately within each column and are therefore intended for within-column comparisons only. **(a)** Reference-based methods, excluding foundation models, were evaluated across classification accuracy, spatial structure preservation, and computational scalability. Each dataset column corresponds to a specific reference–query evaluation setting. MERFISH denotes the internal spatial reference setting, in which 17 MERFISH sections were used as the reference and one section was used as the query. MERFISH (single nucleus) denotes the setting in which a matched single-nucleus RNA-seq reference was used to annotate the same MERFISH query section. Open-ST denotes the two-section setting, with one section used as the reference and the other as the query. Xenium denotes the single-nucleus RNA-seq reference setting, in which twelve spatial sections were evaluated separately as query sections and the resulting metric values were averaged across sections. Stereo-seq (regeneration) denotes the matched-stage regeneration setting, in which, among three sections collected at 5 days post-injury, two sections were used as the reference to annotate the remaining section. Stereo-seq (developmental) denotes the cross-stage developmental setting, in which the later developmental stage 54 was used as the reference to annotate the earlier developmental stage 44. **(b)** Reference-free methods were evaluated using structureaware metrics and scalability metrics.

Across all three dimensions, scANVI and TACCO emerged as the most consistently well-balanced methods. Both achieved high classification accuracy across multiple datasets, ranked highly on structure-aware metrics, suggesting that accurate cell type assignments were accompanied by preservation of biologically meaningful spatial organization, and operated with modest computational requirements (Fig. 2a). In contrast, the performance of many other methods varied substantially across datasets. For example, GraphST achieved competitive performance on the Stereo-seq developmental axolotl brain dataset but performed markedly worse on the remaining datasets. Moreover, GraphST exhausted available system memory when processing the kidney dataset, highlighting potential scalability challenges for large or high-density spatial transcriptomics datasets. Unsupervised approaches, including BANKSY, SpaGCN, and STdeconvolve, exhibited particularly pronounced dataset-dependent behavior. SpaGCN showed relatively strong spatial clustering performance on the Stereo-seq datasets, particularly in AMI, ARI, and AvgBIO, whereas STdeconvolve performed competitively on the Open-ST dataset while achieving the lowest execution time and peak memory consumption (Fig. 2b).

Together, these results indicate that method performance reflects a trade-off among predictive accuracy, spatial structure preservation, and computational efficiency, rather than a single universally optimal solution. We next carefully examined each dataset to determine how tissue context, spatial technology, reference-query settings and biological dynamics influenced annotation performance.

### Benchmarking spatial cell type annotation across reference-transfer settings

The MERFISH mouse spinal cord dataset, recently generated by our group, comprises 18 spatial sections spanning the cervical, thoracic, lumbar, and sacral regions [24]. Each section was profiled using MERFISH with a custom-designed panel of 500 genes, capturing 60 distinct cell types that encompass major neuronal and non-neuronal cell populations across the spinal cord. Specifically, these overarching categories include diverse non-neuronal lineages such as astrocytes, oligodendrocytes, oligodendrocyte progenitors and oligodendrocyte precursor cells (OPCs), microglia, endothelial cells, meningeal cells, ependymal cells, and peripheral glial cells, as well as highly resolved neuronal subtypes, including dorsal excitatory neurons, medial-ventral excitatory neurons, dorsal inhibitory neurons, medial-ventral inhibitory neurons, motor neurons (MN) and cholinergic interneurons. The number of segmented cells varies substantially across anatomical regions, with cervical sections containing the largest number of cells (approximately 6,000 cells per section), whereas sacral sections contain markedly fewer cells (approximately 300 cells per section), reflecting region-specific differences in tissue size and cellular composition. One cervical section was held out as the test dataset, while the remaining 17 sections were used as reference for internal label transfer. Reference sections were incrementally incorporated to progressively increase the size and diversity of the reference set. Annotation performance was evaluated using a multi-faceted benchmarking framework that included quantitative accuracy metrics, cross-reference label consistency, and biological interpretability.

For the selected cervical section, annotation performance generally improved as additional MERFISH reference sections were incorporated, particularly for transfer learning–based methods such as scANVI and Spatial-ID (Fig. 3a). With all 17 reference sections, Spatial-ID achieved the highest overall accuracy (0.792), followed closely by scANVI (0.790), Seurat (0.763), and TACCO (0.738), indicating strong cell-level annotation accuracy for these methods.

**Figure 3:**
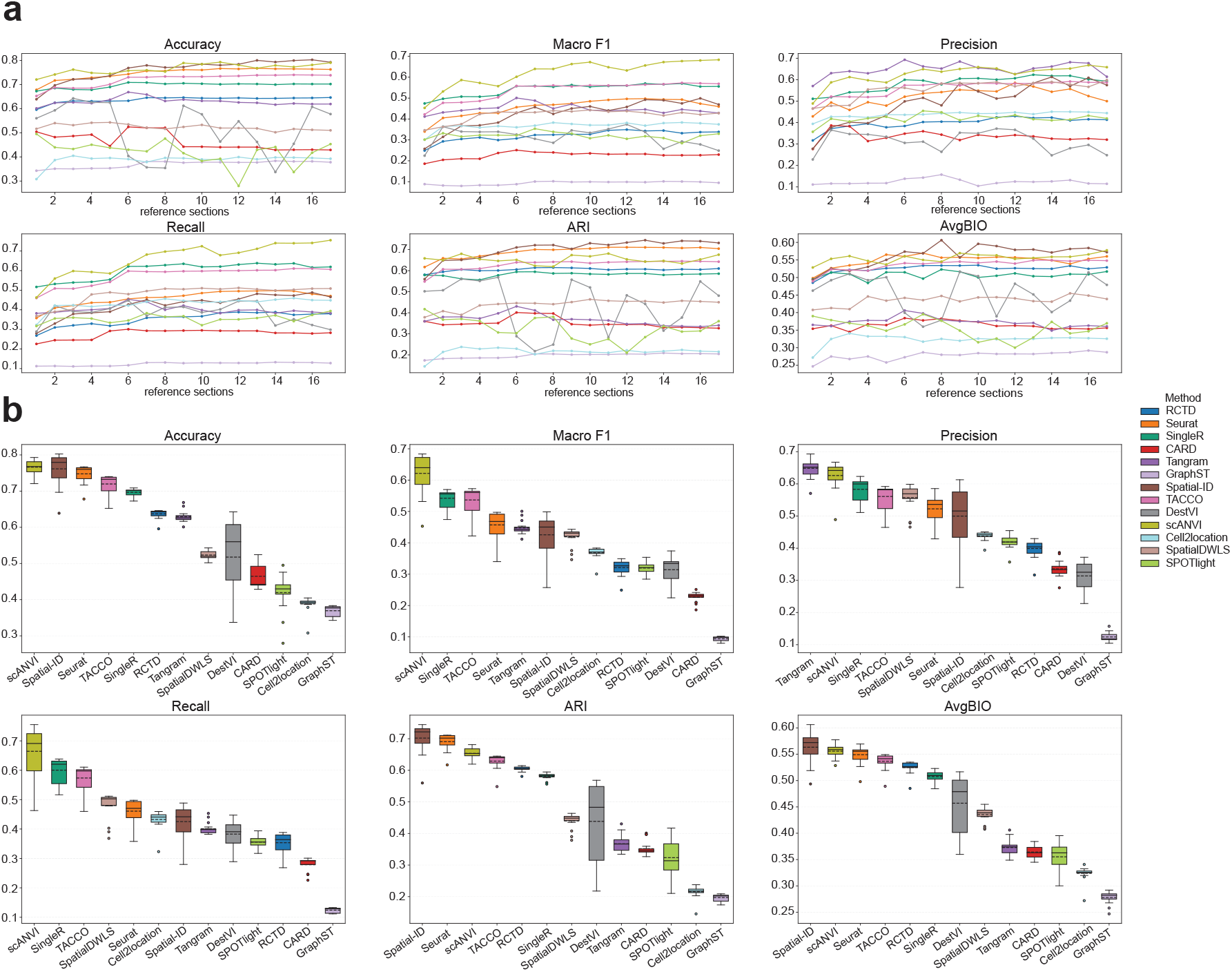
Performance of annotation methods with increasing reference MERFISH volume. **(a)** Line plots showing method performance as the number of reference MERFISH sections increases. Each curve represents one method, and panels report Accuracy, Macro F1, Precision, Recall, ARI, and AvgBIO across reference set sizes. **(b)** Distribution of method performance across all reference set sizes shown in **(a)**. For each method, values from all reference-section-number settings were summarized as box plots for each metric, providing an overview of both average performance and sensitivity to reference volume.

Since overall accuracy can be dominated by abundant cell populations, we further evaluated balanced classification metrics, including macro F1, precision, and recall. scANVI consistently achieved the highest macro F1 across nearly all reference-section settings (Fig. 3a-b), indicating superior performance in balancing annotation accuracy across both abundant and underrepresented cell populations. This observation was further supported by its leading recall, which increased steadily with additional reference sections, suggesting robust recovery of true cell identities across all expanding reference label space. Tangram achieved the highest precision across most settings, indicating highly specific label assignments. However, its lower ARI and AvgBIO indicate that these accurate local predictions did not translate into equally strong preservation of overall label consistency or biological organization. Thus, Tangram may favor conservative, high-confidence assignments for subsets of cells, whereas scANVI provides broader and more balanced recovery of cell type diversity. Spatial-ID achieved the highest overall accuracy and ranked among the top-performing methods in ARI and AvgBIO, but its macro F1, precision, and recall were lower than those of scANVI (Fig. 3a-b). This suggests that Spatial-ID is particularly effective at capturing dominant cell type patterns that contribute to overall classification performance, while scANVI provides a more balanced representation of cell type diversity, including rare and fine-grained populations.

Not all methods benefited monotonically from additional reference sections. DestVI, and similarly SPOTlight, showed fluctuating performance across reference-set sizes rather than consistent improvement. DestVI reached its highest annotation accuracy with three reference sections (0.643) but declined with larger reference sets. A similar non-monotonic pattern was observed for ARI, suggesting that the instability extended beyond cell-wise label assignment to the preservation of global annotation structure relative to the ground truth. These findings indicate that increasing the number of reference sections does not universally enhance annotation performance. In deconvolution-based or generative frameworks, larger reference sets may broaden the represented cellular-state landscape, but can simultaneously introduce increased cellular heterogeneity, anatomical variability, and section-specific batch effects. Such complexity may perturb the learned latent representation and consequently reduce the robustness of downstream label transfer. Collectively, these results suggest that the biological and technical composition of the reference data, rather than reference size alone, is a critical determinant of annotation performance.

Beyond classification accuracy, ARI and AvgBIO further suggested that Spatial-ID, scANVI, and Seurat more effectively preserved overall label consistency and biological organization. In contrast, GraphST, Cell2location, CARD, and SPOTlight exhibited lower ARI and AvgBIO scores, indicating weaker agreement with ground-truth annotations and reduced preservation of biologically coherent structure in this MERFISH dataset.

To assess the robustness of spatial cell type annotation methods under different reference-transfer regimes, we compared performance on the same MERFISH query dataset using either an internal reference derived from the same MERFISH study (within-platform reference) or an external reference generated from single-nucleus RNA-seq data [35] (cross-platform reference). Under the within-platform setting, where the reference and query shared the same technology and experimental protocol, Spatial-ID achieved the highest classification accuracy and the strongest clustering agreement as measured by ARI, demonstrating superior preservation of agreement between predicted and ground-truth label partitions when platform-induced domain shift was minimized. In contrast, method rankings changed substantially under the cross-platform transfer setting. RCTD achieved the highest overall accuracy and the strongest Macro F1 score, indicating more balanced performance across cell types while maintaining high sample-level prediction accuracy (Fig. 2a). Beyond aggregate scores, methods exhibited distinct precision–recall trade-offs: RCTD yielded the highest recall, consistent with broader recovery of query cell types, whereas TACCO achieved the highest precision alongside the best ARI, suggesting more conservative label assignments and improved preservation of global annotation structure at the cluster-partition level. Notably, DestVI obtained the highest AMI and AvgBIO, indicating stronger agreement with the reference partition after chance correction and improved preservation of biologically meaningful organization. Collectively, these results demonstrate that annotation performance is highly dependent on the reference-transfer regime and that method rankings are not consistently preserved across within-platform and cross-platform settings.

### Subtype-level benchmarking reveals challenges in resolving fine-grained cellular identities

To assess model performance at a fine-grained resolution, we quantified classification accuracy across all 60 annotated cell subtypes in the MERFISH mouse spinal cord dataset and additionally summarized performance after grouping subtypes into broader biological categories (Fig. 4). We first examined the hierarchical accuracy tree for Spatial-ID, which achieved the highest overall cell-level accuracy in the selected cervical MERFISH section (Fig. 4a). This hierarchical annotation framework was recently curated by our group to capture the extensive transcriptional diversity of spinal cord neuronal populations, which comprise the largest number of annotated cell subtypes [24]. To enable comparison across annotation methods, hierarchical accuracy trees for all remaining methods are provided in Supplementary Fig. 1. Spatial-ID accurately identified several major neuronal classes, including cholinergic, dorsal inhibitory, dorsal excitatory, and intermediate and ventral (M+MV) populations. However, accuracy varied substantially among downstream subtypes. For example, within the Dorsal Maf lineage, classification performance declined at successive levels of the hierarchy, with particularly low accuracy for DH-ex-Rreb1 and DH-ex-Maf/Cpne4 cells. These results indicate that strong overall performance can obscure substantial variability in subtype-level resolution.

**Figure 4:**
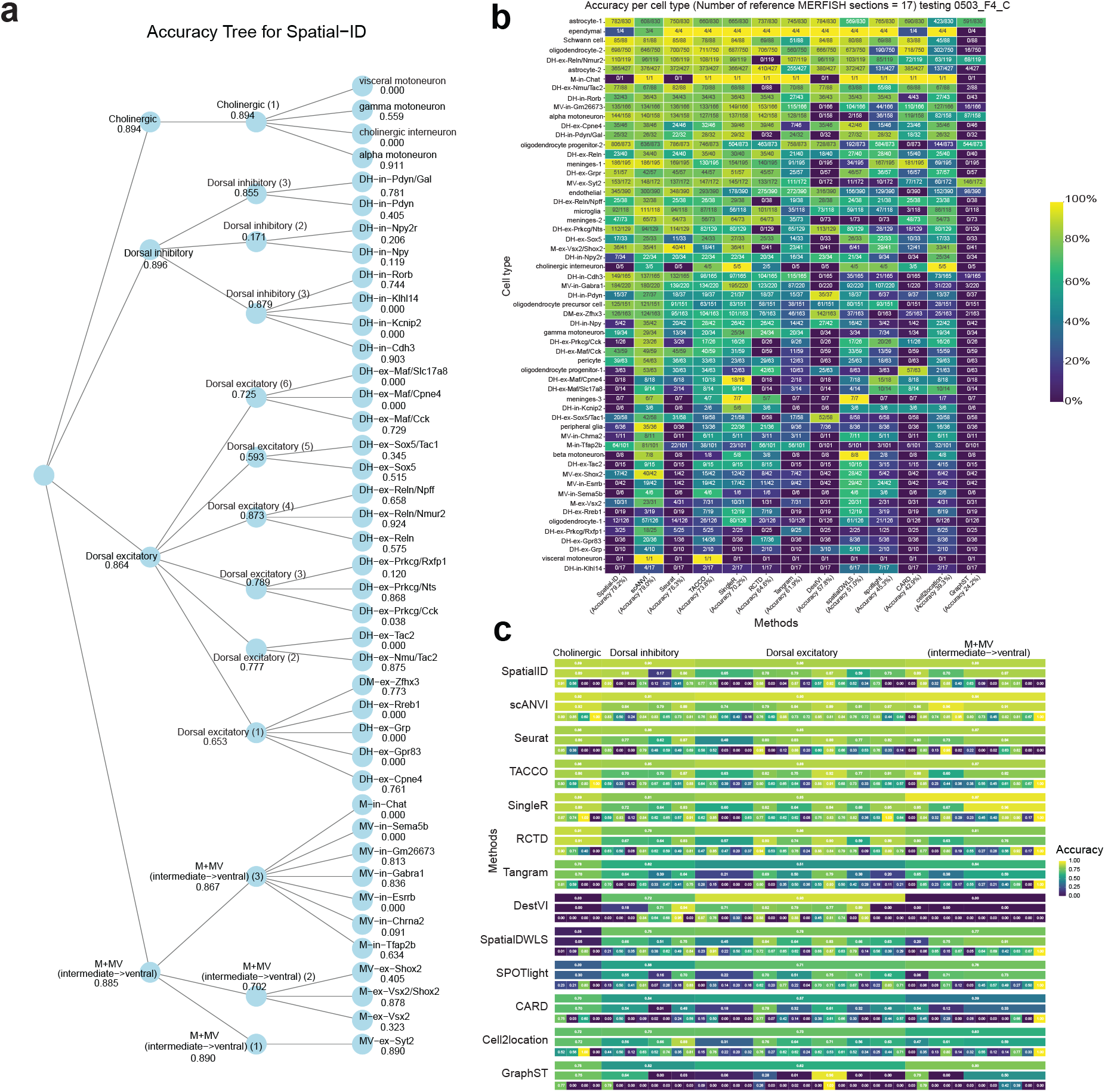
Evaluation of cell type annotation performance across the cell type hierarchy in the MERFISH mouse spinal cord. **(a)** Neuronal cell type hierarchy and classification accuracy produced by Spatial-ID. Cell identities are organized hierarchically from broad neuronal classes to increasingly finegrained subtypes. Internal nodes represent intermediate cell type categories, whereas leaf nodes represent terminal cell types. Values shown beside each node indicate the classification accuracy by Spatial-ID at that level of the hierarchy. **(b)** Heatmap showing annotation accuracy for each cell type across methods in one cervical section of MERFISH mouse spinal cord, using 17 sections as the reference. Rows correspond to cell types and columns correspond to annotation methods. Color intensity indicates the classification accuracy for each cell type using each method. Numbers within each cell indicate the number of correctly annotated cells over the total number of cells for that cell type. This panel highlights differences in method performance at fine cell type resolution. **(c)** Based on the hierarchical classification tree in **(a)**, accuracy was computed for each annotation method at individual classification nodes. Rows are grouped by method, and within each method, separate tracks represent different hierarchical levels or branch-specific classification steps. Columns correspond to nodes in the classification tree. Colors and numeric labels indicate the corresponding accuracy values. This panel summarizes how consistently each method preserves hierarchical cell identity structure across coarse and fine-grained annotation levels.

Interpretation of subtype specific accuracy requires consideration of tissue composition. Certain cell types are sparsely represented or absent in the query cervical spinal cord, such that low classification accuracy may reflect biological absence rather than annotation errors. For example, visceral motoneurons are rare in the cervical regions, and consequently near-zero accuracy in the cervical section is expected (Fig. 4a). In addition, several poorly classified populations corresponded to rare neuronal subtypes, including DHex-Rreb1, which comprises approximately 2% of excitatory neurons in the spinal cord [24]. Together, these observations suggest that subtype abundance contributes substantially to differences in annotation performance.

We next compared subtype-level accuracy across all evaluated methods using the 17-section MERFISH reference setting (Fig. 4b). Among all methods, scANVI achieved the highest mean accuracy across cell subtypes, indicating robust performance in resolving fine-grained cellular identities. Notably, scANVI maintained relatively high accuracy for several rare populations, including meninges-3, MV-in-CHRNA2, beta motoneurons, and MV-in-Sema5b, which were classified with substantially lower accuracy by most other methods. In contrast, Spatial-ID achieved the highest overall cell-level accuracy, but exhibited greater variability across subtypes, consistent with its lower macro F1 score and the patterns observed in the hierarchical annotation tree. Across methods, non-neuronal populations, including astrocyte-1, Schwann cells, and oligodendrocyte-2, were classified with consistently high accuracy, potentially reflecting their transcriptional distinctiveness and relatively large representation in the dataset. By contrast, many neuronal subtypes, particularly finegrained dorsal excitatory and inhibitory populations, exhibited lower and more heterogeneous performance across most methods, reflecting the challenge of distinguishing closely related neuronal identities.

Finally, to determine whether classification errors were concentrated within specific biological lineages, we grouped subtypes into major cell type categories and evaluated category-level performance (Fig. 4c). This analysis showed that performance was highly lineage-dependent. Most methods achieved relatively high accuracy for broader cholinergic, dorsal inhibitory, dorsal excitatory, and M+MV categories, although their ability to resolve fine subtypes within these groups differed markedly. scANVI and TACCO showed comparatively more stable performance across major categories, whereas methods such as Tangram, DestVI, SpatialDWLS, SPOTlight, and GraphST displayed more pronounced lineagespecific reductions in accuracy. Collectively, these results reveal a substantial variation in annotation performance across cellular hierarchies, suggesting that subtype resolution remains influenced by lineagespecific transcriptional similarity and cell type abundance.

### Benchmarking reveals distinct trade-offs between annotation accuracy and biological fidelity

Open-ST is a sequencing-based spatial transcriptomics platform that enables untargeted, whole-transcriptome profiling at single-cell resolution. We analyzed an Open-ST dataset of the human metastatic lymph node using two annotated sections with 38,555 and 39,167 cells, respectively [25]. The human metastatic lymph node dataset provides a stringent benchmark for spatial cell type annotation, capturing dense immune, stromal, and tumor populations with pronounced spatial heterogeneity and complex tissue architecture that were independently validated by histopathology and imaging-based spatial transcriptomics. One section was designated as the reference dataset and the other as the query dataset for benchmarking cell type annotation performance across methods. Quantitative benchmarking revealed that scGPT (partial fine tuning) achieved the highest classification accuracy (0.672) (Fig. 5a). Among non–foundation-model approaches, scANVI (0.655) and Spatial-ID (0.644) ranked among the top-performing methods and also achieved high macro F1. scCello (partial fine tuning) also achieved a competitive accuracy of 0.632, further demonstrating the effectiveness of foundation-model-based approaches in this dataset. In contrast, reference mapping and head tuning strategies yielded comparatively lower accuracy, with scGPT (reference mapping) reaching 0.570, scGPT (head tuning) reaching 0.467, and scCello (head tuning) reaching 0.585.

**Figure 5:**
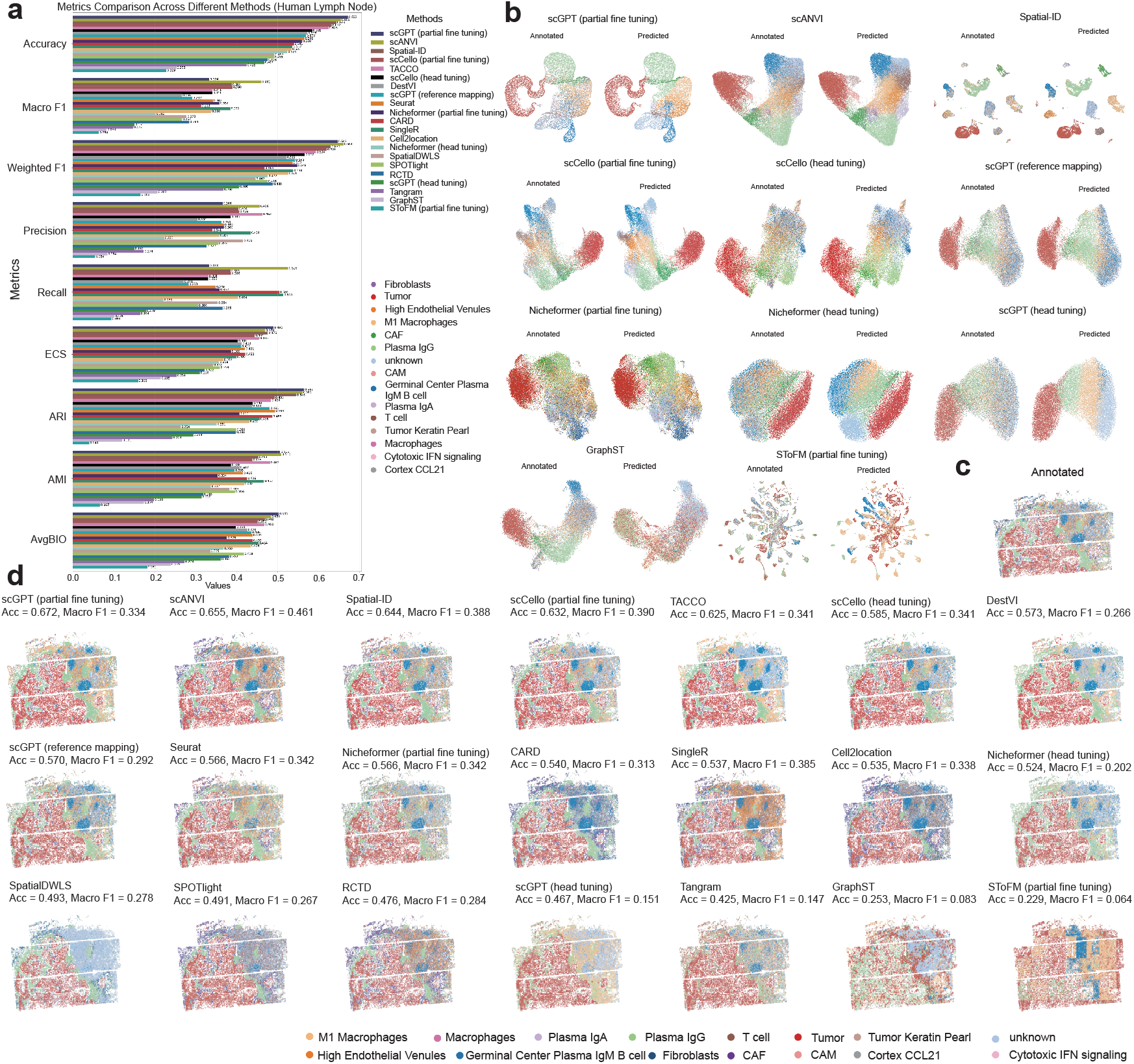
Evaluation of cell type annotation methods using quantitative metrics, latent embeddings, and spatial label distributions in the human lymph node dataset. **(a)** Quantitative comparison of annotation performance across methods in the human lymph node dataset. Bar plots summarize multiple evaluation metrics, including Accuracy, Macro F1, Weighted F1, Precision, Recall, ECS, ARI, AMI, and AvgBIO. Methods are ranked within each metric according to Accuracy. **(b)** Latent representation embeddings for methods that provide low-dimensional cell embeddings. For each method, the embedding is shown with cells colored by ground-truth annotation (“Annotated”) or by the corresponding method-predicted labels (“Predicted”), allowing visual assessment of how well the learned latent space preserves cell type structure and aligns with the annotation results. **(c)** Spatial distribution of the ground-truth cell type annotation in the human lymph node section. Cells are colored according to the reference annotation and shown in the original tissue coordinates. **(d)** For each method, predicted cell type identities were mapped to the original tissue coordinates of the human lymph node section. A shared color scheme was used for all panels to facilitate cross-method comparison. Accuracy (Acc) and macro F1 score are reported above each panel for each method. This visualization highlights differences in the ability of annotation methods to recover spatially coherent cell type distributions.

To qualitatively assess annotation consistency, we visualized predicted cell type labels in both the spatial domain and the latent representation space (Fig. 5b-d). Specifically, predicted labels were projected onto tissue coordinates to examine spatial coherence, while latent embeddings were visualized for methods that explicitly learn low-dimensional representations. For methods that do not optimize a latent embedding, evaluation was restricted to spatial visualizations and quantitative metrics. Our benchmarks demonstrated that both head tuning and reference mapping strategies using foundation models (e.g., scGPT and scCello) generated spatially coherent predictions (Fig. 5d), suggesting that pretrained representations capture substantial biological structure and spatial organization even under limited supervision. However, the transition from head tuning to partial fine tuning yielded a marked gain in resolving micro-heterogeneity. While most methods recovered broad tissue architecture, substantial differences emerged in the spatial coherence and local continuity of their predictions. Head-tuned foundation models, including scGPT, scCello, and Nicheformer, generally recovered broader spatial organization but frequently exhibited diffuse boundaries between adjacent niches. In contrast, scGPT (partial fine tuning) produced better-separated cell populations in the representation space and more spatially coherent domains in tissue space, suggesting improved resolution of microenvironmental organization within the metastatic lymph node (Fig. 5b,d). Notably, strong label-agreement metrics did not always translate into spatially coherent predictions. For instance, Nicheformer (partial fine tuning) achieved moderate cell-level accuracy but a comparatively lower ECS, suggesting that accurate label assignment does not necessarily imply preservation of local spatial neighborhoods (Supplementary Fig. 2).

To determine whether annotation methods preserved the transcriptional programs of tumor microenvironment (TME) populations beyond categorical label agreement, we selected four representative cell types: tumor cells, cancer-associated fibroblasts (CAFs), fibroblasts, and Plasma IgG cells (Fig. 6a). These populations span malignant, stromal, and adaptive immune compartments of the TME and possess well-established marker programs, enabling pathway scoring using single-cell ssGSEA. We compared the enrichment of the top 10 Ground Truth pathways across annotation methods for each target cell type (Fig. 6b). Pathway-level analyses revealed cell type dependent differences in the preservation of biological programs. Tumor cells, CAFs, and Plasma IgG cells exhibited relatively consistent pathway enrichment patterns across methods, indicating that most approaches recovered their core transcriptional programs. In contrast, fibroblasts showed substantially greater variability in pathway enrichment scores, suggesting increased sensitivity to the choice of annotation method. This observation is consistent with the greater transcriptional heterogeneity of stromal populations and highlights differences in the ability of annotation methods to recover biologically coherent cellular states. Collectively, these results demonstrate that annotation performance extends beyond label assignment accuracy and should also be evaluated by the extent to which annotated populations retain their expected transcriptional programs.

**Figure 6:**
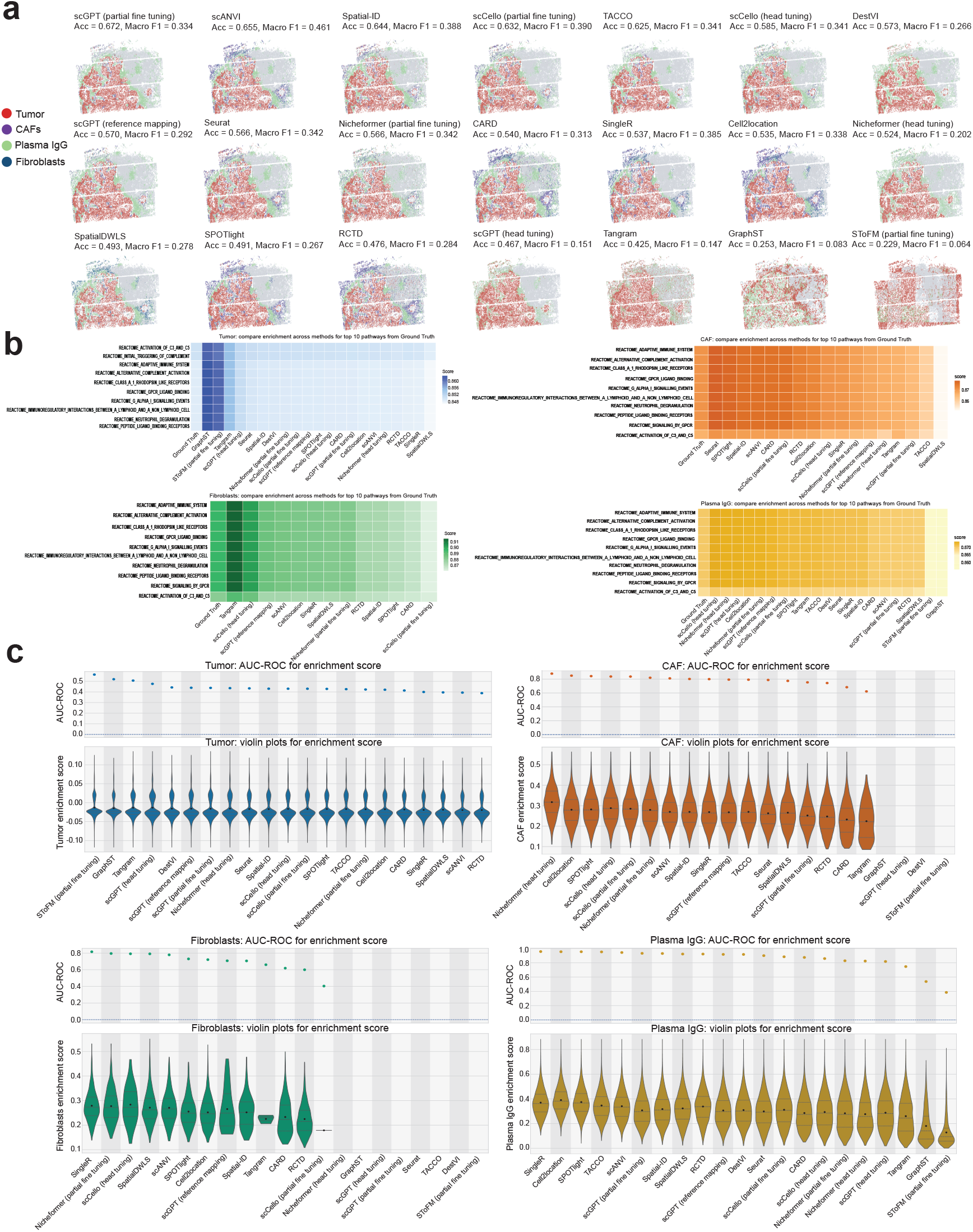
Comparison of cell type annotation methods, enrichment-based label concordance, and downstream pathway consistency in the human lymph node dataset. **(a)** Spatial visualization of representative cell type annotations across methods. Four representative cell types (Tumor cells, CAF, Fibroblasts, and Plasma IgG cells) were visualized in tissue coordinates for the human lymph node dataset to compare annotation performance across methods. For each method, cells assigned to the selected cell types are highlighted in distinct colors, while all remaining cells are shown in gray. Annotation accuracy (Acc) and macro F1 score are reported above each panel. **(b)** Comparison of downstream pathway enrichment across methods for representative cell types. For each representative cell type, the top 10 pathways were first identified from the ground truth annotation and are shown as the first column in each heatmap. The corresponding pathway enrichment patterns were then compared across annotation methods using the same pathway set identified from ground truth annotation. **(c)** For each representative cell type, we computed the AUC-ROC between the method-derived binary labels and the ground-truth enrichment scores. In each cell type panel, the upper scatter plot shows the AUC-ROC for each method, whereas the lower violin plot shows the distribution of ground-truth enrichment scores for cells grouped according to the method-assigned labels.

In addition to pathway-level comparisons, we quantified the concordance between method-derived annotations and cell type specific transcriptional programs using the area under the receiver operating characteristic curve (AUC-ROC) based on ssGSEA scores (Fig. 6c). Program concordance was strongly cell type dependent, with no single method consistently achieving the highest AUC-ROC across all populations. Instead, distinct methods performed best for different cellular compartments, including SToFM (partial fine tuning) for tumor cells, Nicheformer (head tuning) for CAFs, and SingleR for both fibroblasts and Plasma IgG cells. Notably, rankings based on program concordance frequently differed from rankings based on classification accuracy. Despite this overarching trade-off, SPOTlight, Nicheformer (partial fine tuning), scANVI, Spatial-ID, SingleR, and Cell2location demonstrated consistent performance across both metrics, reflecting a more uniform alignment with native biological programs across diverse cell types. These results demonstrate that label assignment accuracy and transcriptional program preservation capture complementary aspects of annotation quality and should be evaluated jointly when assessing spatial cell type annotation performance.

### Benchmarking highlights challenges in annotating dynamic cellular states during kidney injury and repair

Xenium is a high-resolution, imaging-based spatial transcriptomics platform from 10x Genomics that profiles RNA expression across tissue sections at single-cell resolution, enabling integrative analysis of cell types and their spatial contexts in complex tissues [36]. We analyzed a Xenium dataset comprising 12 mouse kidney samples across six stages of injury and repair (Sham, Hour 4, Hour 12, Day 2, Day 14, and Week 6 post-injury), with two technical replicates for each time point and a targeted panel of 300 genes [2]. This dataset was linked to a matched singlenucleus RNA-seq reference from the same ischemiareperfusion injury (IRI) time course, which informed the targeted panel design and supported reference-based validation of Xenium cell type annotations [37]. It therefore provides a stringent benchmark for cell type annotation because acute injury induces substantial transcriptional remodeling and generates transient cellular states that are difficult to map to static reference atlases.

To evaluate the biological concordance between method-derived annotations and manual reference labels, we computed differential marker genes separately for the reference annotations and for each method-derived cell type group, and quantified the overlap among the top 10, 20, and 30 ranked markers (Fig. 7a). Overall, most methods exhibited relatively stable marker recovery across different marker cutoffs, indicating that the highest-ranking cell-typespecific markers were generally preserved regardless of the number of differential genes considered. Among all methods, scANVI, TACCO, SingleR, and Seurat consistently achieved the highest scores across sections, indicating superior recovery of reference marker-gene signatures. In contrast, CARD showed substantially lower overlap scores across all sections, indicating limited agreement with the marker structure defined by the manual annotations. For most methods, overlap scores were highest when considering the top 10 differential genes and declined only modestly when extending the comparison to the top 20 or 30 genes, indicating that marker-gene recovery was generally robust across different ranking thresholds.

**Figure 7:**
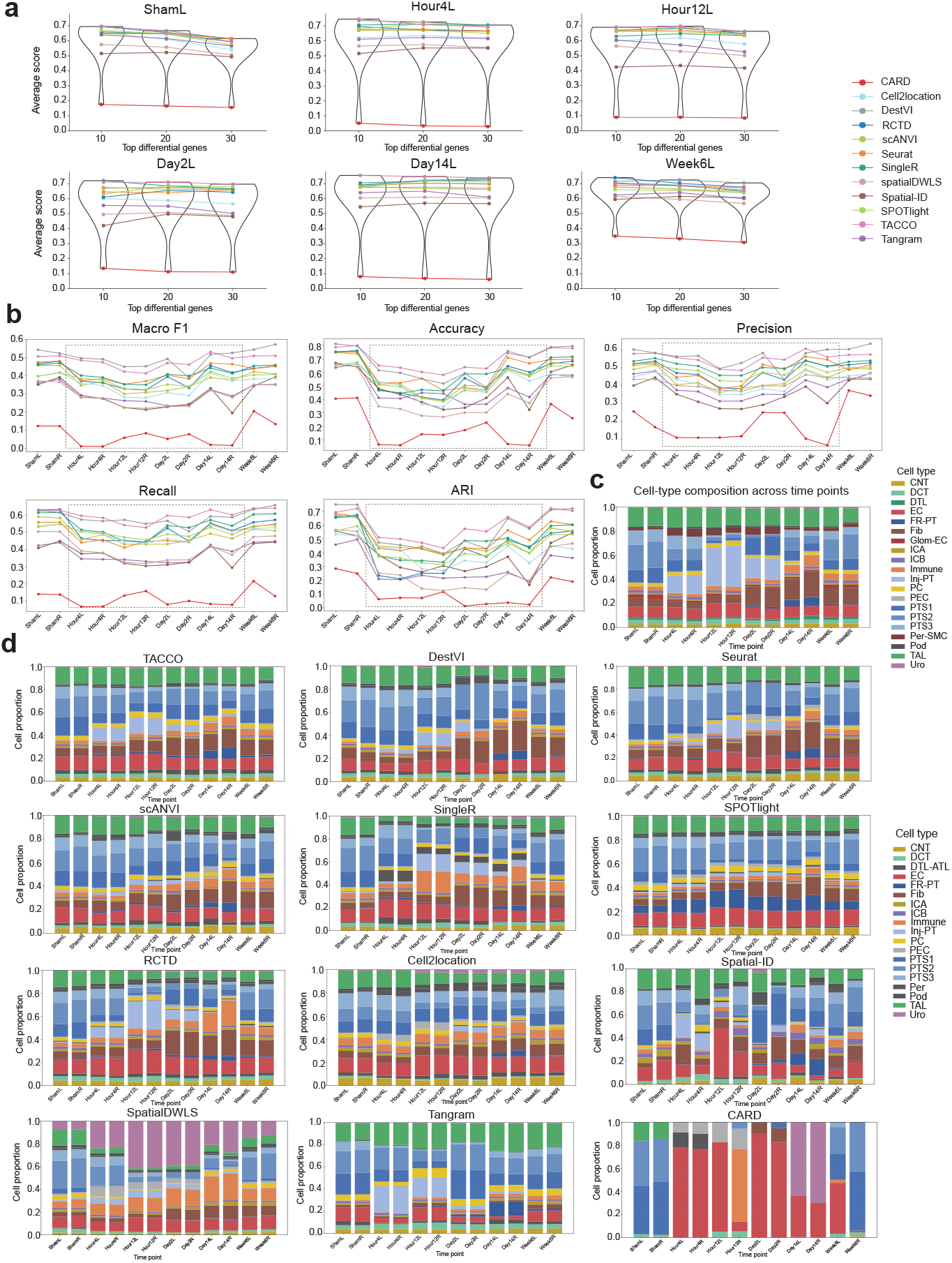
Comparative evaluation of cell type annotation methods in the kidney injury and repair dataset. **(a)** Marker gene recovery analysis. For each method, differential genes identified in each section were ranked and compared with reference differential genes derived from the ground-truth annotations. Performance was summarized as the average overlap score between the top-ranked method-derived genes and the corresponding reference genes for each cell type, using the top 10, 20, and 30 differential genes. **(b)** Temporal dynamics of annotation performance during kidney injury and repair. Because each tissue section corresponds to a specific time point, performance metrics are shown for individual sections along the time course. Macro F1, accuracy, precision, recall, and adjusted Rand index (ARI) were used to evaluate concordance between predicted cell type labels and ground-truth annotations. Dashed lines mark transitions between major biological stages of the kidney injury–repair process, highlighting the acute injury interval and subsequent repair phases during which annotation performance varied substantially. **(c)** Temporal changes in cell type composition for ground truth annotation. Stacked bar plots display the relative abundance of cell types across sections collected at different time points. Pod, podocytes; Glom-EC, glomerulus endothelial; PTS1, S1 segment of proximal tubule; PTS2, S2 segment of proximal tubule; PTS3, S3 segment of proximal tubule; Inj-PT, injured proximal tubule; FR-PT, failed repair proximal tubule; DTL, descending limb of loop of Henle; TAL, thick ascending limb of loop of Henle; DCT, distal convoluted tubule; CNT, connecting tubule; PC, principal cells; ICA, type A intercalated cells; ICB, type B intercalated cells; Uro, urothelium; PEC, parietal epithelial cells; EC, endothelial cells; Fib, fibroblasts; Per-SMC, pericytes and smooth muscle cells. **(d)** Method-specific estimates of cell type proportion across kidney injury-and-repair time points.

Current cell type annotation methods are fundamentally challenged under dynamic pathological conditions, where cells frequently occupy transitional or hybrid states that are not adequately represented by discrete reference-defined cell types. Consistent with these biological complexities, we observed substantial performance variability across disease stages, with annotation accuracy generally declining during acute injury phases and partially recovering during later remodeling stages (Fig. 7b). Xenium datasets collected along injury–repair time courses pose particular challenges for cell type annotation due to the high prevalence of transitional and maladaptive cellular states. The largest performance declines were observed during the acute injury interval (Hour 4 – Day 2). Despite these difficulties, TACCO and DestVI consistently ranked among the top-performing methods, maintaining relatively high Macro F1, accuracy, precision, recall, and ARI across the kidney injury–repair time course.

Because changes in cell type composition reflect underlying pathological processes of kidney injury and repair, we further compared the cell type proportions predicted by each method across disease stages (Fig. 7d). The ground-truth compositions delineated a stereotyped injury-and-repair trajectory: the injured proximal tubule (Inj-PT) population transiently expanded during the acute injury phase, reaching its highest proportions at Hour 4, Hour 12, and Day 2 before contracting as the tissue progressed into repair, while a smaller failed-repair PT (FR-PT) population emerged at the later time points (Day 14, Week 6) (Fig. 7c). Compared with these biologically interpretable trajectories, TACCO most faithfully recapitulated the temporal dynamics observed in the reference annotations, accurately capturing both the transient expansion of Inj-PT cells during acute injury and the emergence of FR-PT cells during the repair phase. DestVI and Seurat also broadly recovered these trends, although with greater smoothing of temporal changes and reduced sensitivity to some injury-associated populations. These results indicate that the underlying biological signal is recoverable from the spatial data when methods are sufficiently flexible to accommodate disease-induced states. However, CARD-derived compositions exhibited pronounced distortions, including abrupt dominance of individual cell types and loss of expected temporal continuity. Furthermore, CARD exhibited limited sensitivity to the rare injury-associated states, yielding negligible signals for Inj-PT and FR-PT across all time points. This failure likely reflects CARD’s heavy reliance on reference-derived proportions and its limited capacity to accommodate transitional or injury-associated cell states, which tend to be collapsed into adjacent normal categories under dynamic conditions, highlighting that recovery of disease-relevant cell states, not only canonical lineages, should be a core criterion when benchmarking cell type annotation methods on disease time courses.

### Dynamic cellular states complicate cell type annotation during regeneration and development

We next evaluated cell type annotation methods in axolotl telencephalon datasets spanning regeneration and development. These datasets provide a biologically challenging setting because cell identities are influenced by both spatial organization and dynamic state transitions. In the regeneration experiment, two sections from 5 days post-injury (5 DPI) were used as references and an independent section from the same stage was used as the query, thereby assessing label transfer within a matched regenerative state. Across methods, SingleR achieved the strongest overall performance, ranking highest in accuracy (0.597), macro F1 (0.405), weighted F1 (0.589), ECS (0.405), ARI (0.438), and AMI (0.553) (Fig. 8a). Seurat and TACCO also performed competitively across multiple metrics, with Seurat showing the highest AvgBIO (0.451), indicating relatively strong preservation of biological structure after label transfer. In contrast, methods such as Tangram, DestVI, and GraphST showed lower overall performance in this setting. Notably, even the best-performing methods achieved only moderate macro F1 and ARI values, indicating persistent difficulty in resolving fine-grained cell identities despite successful recovery of broad cellular organization. This observation is consistent with the biology of axolotl brain regeneration, where injury-induced ependymoglial activation and transitional progenitor and immature neuronal states generate a continuum of cellular identities during neuronal replenishment, thereby blurring discrete cell type boundaries.

**Figure 8:**
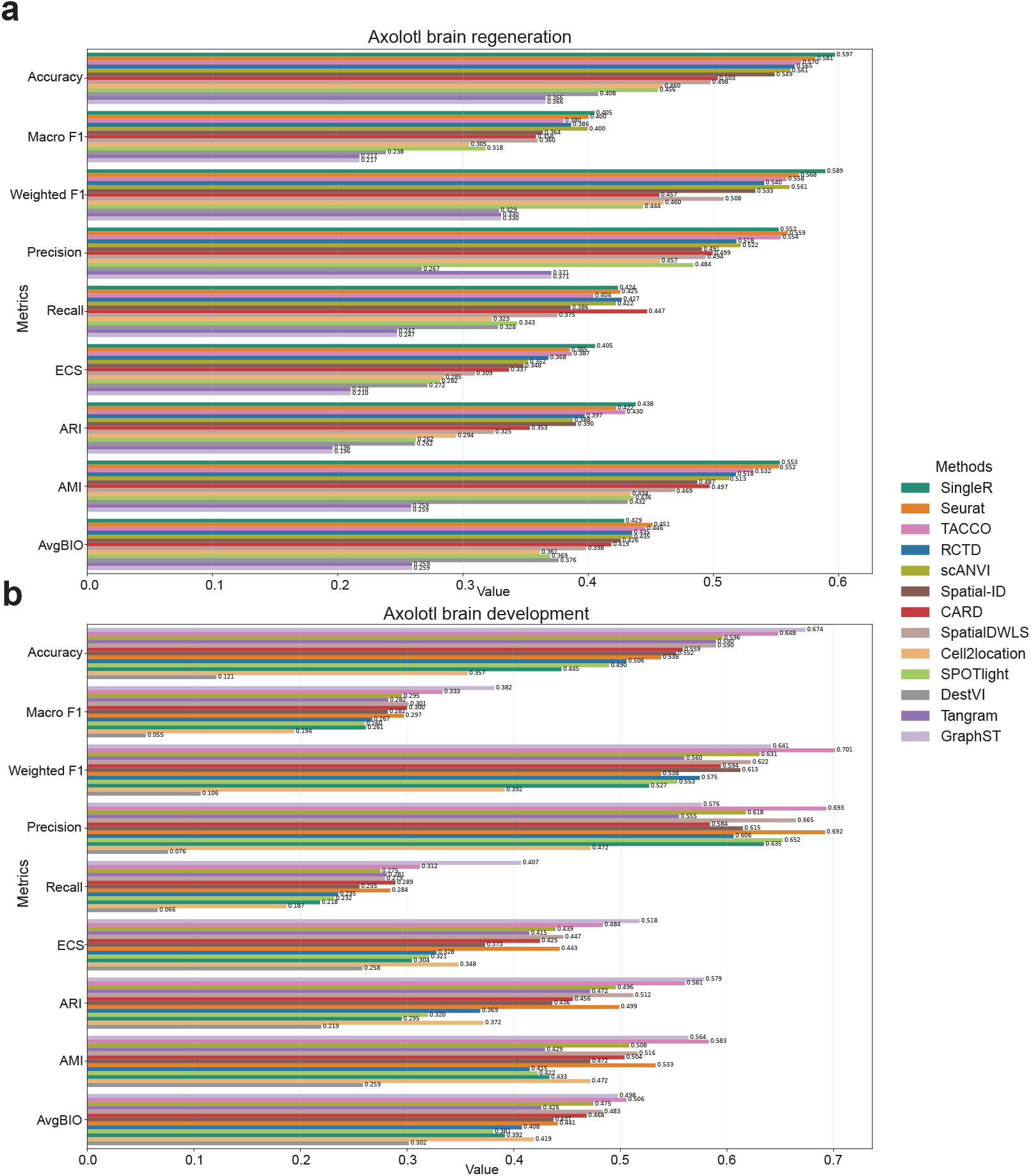
Benchmarking cell type annotation methods on the Stereo-seq axolotl brain dataset under within-stage and cross-stage label transfer settings. **(a)** Performance of 13 cell type annotation methods on the axolotl brain regeneration dataset (within-stage label transfer). Three biological replicate slices from the same developmental stage were used, with two slices serving as the reference and the remaining slice used as the query. **(b)** Performance of the same 13 methods on the axolotl brain development dataset (cross-stage label transfer), in which one stage 54 slice was used as the reference and one stage 44 slice as the query. Performance was evaluated using nine metrics, including classification accuracy (Accuracy, Macro F1, Weighted F1, Precision, and Recall), clustering agreement with ground-truth labels (ARI, AMI, and ECS), and an aggregated biological-conservation score (AvgBIO). Exact values are shown at the end of each bar with higher values indicating better performance for all metrics.

We further evaluated label transfer across distinct developmental stages, using stage 54 sections as the reference and a stage 44 section as the query. In this setting, one query-specific population, dNBL3, was absent from the stage 54 reference annotation. Because most supervised label-transfer methods operate within a closed label space defined by the reference, cells belonging to a reference-absent population cannot be assigned their true identity and are instead forced into the most similar available reference category. We therefore excluded dNBL3 cells from calculations of accuracy, macro F1, weighted F1, precision, recall, ECS, ARI, AMI, and AvgBIO. This exclusion was applied uniformly across all benchmarked methods and prevented artificial penalization arising from an unavailable reference label. Nevertheless, the presence of dNBL3 highlights an important limitation of current reference-based annotation frameworks in developmental systems: stage-specific or previously unseen cell states are typically misclassified rather than explicitly identified as novel or out-of-distribution populations.

Method rankings differed substantially from those observed in the regeneration benchmark. GraphST emerged as the strongest performer on balanced evaluation metrics, achieving the highest accuracy (0.674), macro F1 (0.382), recall (0.407), ECS (0.518), and ARI (0.579) (Fig. 8b). These results suggest that incorporation of spatial graph information may confer an advantage when transferring labels across developmental stages characterized by substantial transcriptional divergence. TACCO remained highly competitive, ranking highest in weighted F1 (0.701), precision (0.693), AMI (0.583), and AvgBIO (0.506), reflecting strong performance on abundant and well-represented cell populations. Seurat also maintained robust performance across most metrics, whereas Cell2location, SPOTlight, DestVI, and Tangram showed substantially degraded performance. Overall, these findings indicate that method performance is highly context dependent: approaches that perform well in within-stage regenerative transfer do not necessarily retain the same advantage in cross-stage developmental transfer, highlighting the importance of benchmarking annotation methods across biologically diverse scenarios.

### LLM-based annotation captures broad biological lineage structure

To assess whether LLMs can recover biologically meaningful cell identities from marker-based descriptions, we benchmarked GPT-4o and Claude Sonnet 4-5 across four spatial transcriptomics datasets spanning neural, renal injury, developmental/regenerative, and tumor-microenvironment contexts. For each dataset, we first performed de novo clustering using Leiden (Fig. 9a-h) and BANKSY (Supplementary Fig. 3), selecting clustering resolution to approximately match the number of groundtruth cell type clusters to facilitate one-to-one comparison with manual annotations. We then prompted each LLM to assign biological identities to the resulting clusters using only the top 10 differentially expressed marker genes. Because LLM-generated labels are unconstrained and may differ semantically from reference annotations, we reordered the rows (ground-truth labels) and columns (predicted labels) of each row-normalized confusion matrix according to biological lineage correspondence. Specifically, for each (predicted, ground-truth) label pair, GPT-4o was used as a semantic evaluator to determine the closest biological correspondence, and both axes were arranged such that biologically matched populations aligned along the diagonal. Under this ordering, accurate annotation is reflected by concentrated diagonal enrichment, whereas off-diagonal signal indicates systematic confusion between biologically distinct or transcriptionally adjacent populations.

**Figure 9:**
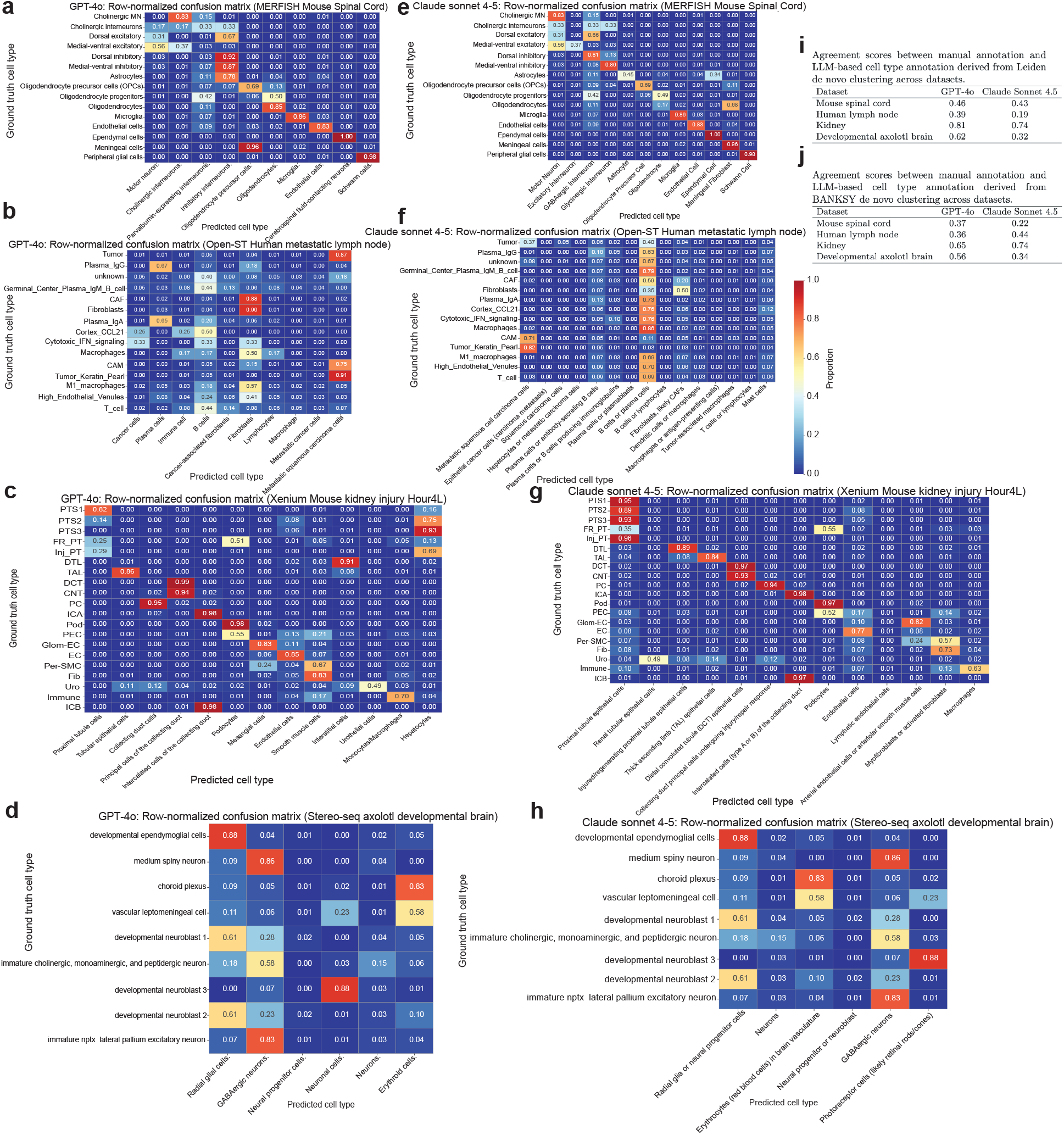
Evaluation of LLM-based cluster annotation across all datasets. **(a–h)** Row-normalized confusion matrices for LLM-based cell type annotation across four spatial transcriptomics datasets. For each dataset, de novo clustering was first performed using Leiden, followed by cluster-level annotation with GPT4o or Claude Sonnet 4-5. Both ground-truth and predicted cell type labels were ordered by biological lineage. **(i–j)** Agreement scores between manual annotations and LLM-derived cell type labels. For each groundtruth/predicted label pair, GPT-4o was prompted to classify the semantic correspondence as a perfect match, partial match, or no match, which were assigned scores of 2, 1, and 0, respectively. Scores were then averaged within each dataset. **(i)** Results based on Leiden-derived clusters. **(j)** Results based on BANKSY-derived clusters. Higher scores indicate stronger semantic concordance between predicted and reference labels.

To complement the visual diagonal pattern with a quantitative measure, we further evaluated label-level semantic agreement using GPT-4o as an evaluator: for each (ground-truth, predicted) label pair, the model was prompted to classify the correspondence as a perfect match (score 2), partial match (score 1), or no match (score 0), and the resulting per-label scores were averaged across cells within each dataset to yield a single agreement score per dataset–LLM combination (Fig. 9i-j) [18]. Higher scores indicate stronger semantic concordance between predicted and groundtruth labels.

Across datasets, both GPT-4o and Claude Sonnet 4-5 recovered broad lineage-level structure, as indicated by enrichment of predictions along or near the diagonal of the confusion matrices rather than diffuse off-diagonal assignments. However, annotation performance was highly context-dependent, with agreement scores varying across datasets and clustering strategies (Fig. 9i-j). The mouse kidney dataset showed the strongest concordance under both Leiden and BANKSY clustering. Using Leiden clusters, GPT-4o and Claude Sonnet 4-5 achieved agreement scores of 0.81 and 0.74, respectively; using BANKSY clusters, the corresponding scores were 0.65 and 0.74. Major epithelial populations, including distal convoluted tubule (DCT), thick ascending limb (TAL), principal cells (PC), type A intercalated cells (ICA), and podocytes (Pod), showed strong diagonal enrichment, with several classes reaching agreement scores of at least 0.85 (Fig. 9c, 9g).

The mouse spinal cord dataset also showed clear lineage-level concordance, particularly for major glial and non-neuronal populations, including oligodendrocytes, microglia, endothelial cells, and peripheral glial cells (Fig. 9a, 9e). In contrast, neuronal subtypes produced more dispersed assignments, consistent with the greater difficulty of resolving closely related neuronal states from marker lists alone. The weakest performance was observed in the human metastatic lymph node dataset (Leiden: GPT-4o, 0.39; Claude Sonnet 4-5, 0.19; BANKSY: GPT4o, 0.36; Claude Sonnet 4-5, 0.44), where predictions were broadly distributed across immune, stromal, and tumor-associated populations. For example, both models frequently collapsed multiple groundtruth populations, including macrophages, CAFs, and plasma-cell subsets, into a generic “B cells” prediction class (Fig. 9b, 9f).

These dataset-specific differences suggest that LLM-based annotation is most reliable when cell identities are defined by canonical and well-separated marker programs, as in many epithelial and major non-neuronal lineages. In contrast, performance declines in tumor microenvironments and developmental or injury contexts, where transcriptional states are continuous, overlapping, and shaped by disease or stage-specific remodeling. Across all four datasets, LLMs generally recovered broad lineage identity but struggled to resolve closely related subtypes and cellular states. For example, in the kidney injury-repair dataset, PTS1, PTS2, and PTS3 were typically recognized as proximal tubule cells, but were frequently collapsed into a single “proximal tubule” class, obscuring segmental subtype distinctions. Similarly, in the developmental axolotl brain dataset, distinct progenitor populations were often merged with radial glia or assigned to a generic “neural progenitor” identity; developmental neuroblast 1 was mapped to radial glial cells in 61% of instances by both LLMs.

Accordingly, most errors occurred between biologically related or developmentally adjacent populations rather than across unrelated lineages, including confusion among proximal tubule subtypes, immature neuronal and progenitor populations, and immune subtypes within shared lymphoid lineages. These results indicate that LLMs capture coarse marker-derived biological semantics but do not reliably distinguish fine-grained, context-dependent cell states. Occasional cross-lineage errors were also observed, such as choroid plexus cells being annotated by both LLMs as “erythroid cells” in the developmental brain dataset (Fig. 9d, 9h) and injured proximal tubule cells being annotated by GPT-4o as “hepatocytes” in the kidney dataset (Fig. 9c). These examples suggest that when canonical markers are incomplete, atypical, or weakly represented, LLMs may assign biologically implausible but lexically plausible identities based on their pretrained knowledge space.

Neither GPT-4o nor Claude Sonnet 4-5 was uniformly superior across settings. Under Leiden clustering, GPT-4o outperformed Claude Sonnet 4-5 in all four datasets, with the largest gaps in the developmental axolotl brain (0.62 versus 0.32) and human lymph node (0.39 versus 0.19). Under BANKSY clustering, however, Claude surpassed GPT-4o in the kidney (0.74 versus 0.65) and lymph node (0.44 versus 0.36) datasets, indicating that model ranking was sensitive to upstream clustering choices. Despite these differences, the overall pattern was consistent across models and clustering methods: agreement was highest in datasets containing well-defined canonical cell types, such as the kidney, and lowest in datasets dominated by heterogeneous, transitional, or pathology-associated cellular states, such as the metastatic lymph node (Fig. 9i-j). Together, these results indicate that general-purpose LLMs can recover broad lineage organization from marker descriptions but are not yet reliable substitutes for dedicated annotation methods when fine discrimination among related, transitional, or pathology-associated cell states is required.

## Discussion

Accurate cell type annotation remains a fundamental prerequisite for interpreting spatial transcriptomics data, yet the increasing diversity of spatial technologies, biological systems, and computational frameworks has made it difficult to determine which annotation strategies are most appropriate for a given study. Through a systematic benchmark spanning four spatial transcriptomics platforms, multiple biological contexts, and twenty state-of-the-art annotation methods, we demonstrate that annotation performance is strongly dependent on biological complexity, reference-query similarity, and the level of cellular resolution being interrogated. Across datasets, no single method uniformly dominated all evaluation scenarios, underscoring the importance of aligning method selection with the biological question, reference availability, and downstream analytical objectives.

Several broad conclusions emerged from this benchmark. First, methods that consistently ranked highly across datasets, including scANVI, TACCO, and Seurat, achieved a favorable balance between classification accuracy, preservation of biological structure, and computational scalability. Importantly, strong performance was not limited to a single methodological paradigm. Rather, successful approaches generally combined robust representation learning with effective mechanisms for integrating reference information while remaining tolerant to technical and biological variability. This observation suggests that robustness to domain shift may be a more important determinant of practical performance than the specific statistical framework underlying a method.

Second, our results reveal that cell type annotation should not be evaluated solely through aggregate accuracy metrics. Across multiple datasets, methods with similar classification accuracy often differed substantially in their ability to preserve higherorder biological organization, recover marker-gene programs, or maintain biologically meaningful pathway activity. In the metastatic lymph node dataset, for example, rankings based on classification accuracy were not always concordant with rankings based on transcriptional-program preservation. These findings indicate that annotation quality encompasses multiple dimensions, including accurate label assignment, preservation of cellular relationships, and maintenance of biologically interpretable gene-expression programs. Consequently, future benchmarking efforts should move beyond cell-level classification metrics and incorporate structure-aware and biology-aware evaluations that better reflect downstream analytical utility.

A major contribution of this study is the systematic assessment of annotation performance across hierarchical levels of cellular identity. While many methods accurately recovered broad lineages, performance declined substantially when resolving closely related subtypes. This pattern was particularly evident in the MERFISH spinal cord dataset, where neuronal subclasses frequently exhibited considerably lower accuracy than broader lineage categories. Such failures likely reflect both biological and technical factors. Closely related neuronal populations often differ by relatively subtle transcriptional programs, exist along continuous functional gradients, and are frequently underrepresented in reference datasets. As a result, conventional label-transfer frameworks may successfully recover lineage identity while failing to distinguish fine-grained cellular states. These findings suggest that reported annotation accuracy may substantially overestimate the ability of current methods to resolve biologically meaningful cellular heterogeneity.

Our analyses further demonstrate that dynamic biological systems represent a particularly challenging setting for cell type annotation. Performance generally declined in datasets characterized by injury, regeneration, or developmental transitions, including the injury-repair kidney model and the axolotl regeneration and developmental datasets. In these contexts, cells frequently occupy intermediate, transitional, or maladaptive states that do not correspond cleanly to discrete reference-defined categories. Because most current annotation frameworks assume a fixed and closed label space, cells exhibiting novel or stage-specific transcriptional programs are often forced into the nearest available reference class. The resulting misclassifications may obscure biologically important cellular trajectories and state transitions. These observations highlight a fundamental limitation of current reference-based annotation paradigms and motivate the development of approaches capable of explicitly identifying out-of-distribution populations, continuous cell-state trajectories, and previously unseen cellular identities.

An important question in spatial transcriptomics is whether explicit incorporation of spatial context improves annotation accuracy. Although spatially informed methods have attracted substantial attention, our benchmark indicates that the benefits of spatial information are neither universal nor consistent across biological settings. Methods that leverage spatial neighborhoods, graph representations, or tissue architecture occasionally provided substantial advantages, particularly in developmental contexts where local spatial organization closely reflects cellular identity. However, spatially aware approaches did not consistently outperform transcription-based methods across all datasets. These findings suggest that the utility of spatial information depends on the extent to which local tissue context is informative for distinguishing cell identities. In tissues characterized by sharp cellular boundaries or well-organized spatial niches, neighborhood information may improve annotation. Conversely, in highly heterogeneous environments containing mixed cellular states or extensive remodeling, spatial proximity alone may provide limited discriminative power. Future methodological development may therefore benefit from adaptive frameworks that dynamically determine when and how spatial context should influence annotation decisions.

The emergence of transcriptome foundation models represents one of the most significant recent developments in computational genomics. Our benchmark demonstrates that foundation-model-based approaches can achieve competitive performance, particularly when pretrained representations are adapted through partial fine tuning. In contrast, head tuning strategies frequently underperformed relative to more extensive adaptation, suggesting that pretrained representations alone may not fully capture the biological variation required for accurate annotation in complex spatial datasets. Although foundation models showed promise, they did not consistently outperform leading reference-based approaches across all benchmark scenarios. These findings indicate that current foundation models provide a strong starting point for annotation but remain limited by challenges including biological domain shift, incomplete representation of rare cell states, and discrepancies between pretraining and target datasets. Continued expansion of pretraining corpora to include diverse spatial, developmental, and disease contexts may improve their generalizability and downstream utility.

Our evaluation of large language models highlights both the promise and current limitations of marker-based annotation paradigms. Across multiple datasets, GPT-4o and Claude Sonnet 4-5 successfully recovered broad lineage-level identities from marker-gene signatures, particularly for canonical epithelial and non-neuronal populations. However, both models struggled to resolve closely related subtypes, transitional states, and pathology-associated populations. Notably, most errors occurred between biologically related cell types rather than unrelated lineages, suggesting that LLMs capture substantial biological semantics despite lacking explicit transcriptomic modeling. Nevertheless, performance remained highly dependent on marker quality, clustering resolution, and biological context. These results support the use of LLMs as efficient tools for preliminary annotation, hypothesis generation, and biological interpretation, while emphasizing that they are not yet substitutes for dedicated quantitative annotation frameworks when high-resolution cell-state discrimination is required.

Several limitations should be considered when interpreting this benchmark. First, although the datasets encompass diverse technologies and biological systems, they cannot capture the full spectrum of tissues, disease states, and spatial transcriptomics platforms currently in use. Second, all benchmark evaluations depend on expert-curated annotations as ground truth, which themselves may contain uncertainty, particularly for transitional or poorly characterized cell states. Third, many methods were originally designed for distinct analytical objectives, including deconvolution, clustering, or label transfer, and may not be optimized for all benchmark settings considered here. Finally, rapidly evolving foundation models and multimodal spatial frameworks may alter the relative performance landscape in the near future.

In summary, our study provides a comprehensive evaluation of contemporary cell type annotation strategies for spatial transcriptomics across multiple technologies, biological contexts, and levels of cellular resolution. We show that fine-grained subtype annotation, dynamic cellular states, and crossdomain transfer remain major challenges for existing methods, while also demonstrating that high classification accuracy alone is insufficient to assess annotation quality. By integrating conventional classification metrics with structure-aware and biology-aware evaluations, this benchmark provides a practical framework for assessing annotation performance and selecting methods appropriate for specific scientific objectives. As spatial transcriptomics continues to expand toward increasingly complex tissues, multimodal measurements, and atlas-scale studies, future annotation frameworks will need to move beyond discrete label assignment toward models capable of representing cellular continua, detecting novel states, and integrating molecular, spatial, and contextual information within a unified framework.

## Methods

### Settings of benchmarked methods

All methods were evaluated using the subset of QC-passed query cells whose ground-truth cell types were represented in the reference dataset. Query cells belonging to cell types absent from the reference were excluded from evaluation.

For algorithms that require explicit specification of the expected number of cell types (e.g., BANKSY), this parameter was set to the true number of cell types defined by the ground truth annotation. For methods that instead rely on a clustering resolution parameter, values were selected according to the official documentation and recommended best-practice guidelines. This approach reflects the performance that users are likely to obtain under realistic application settings. To ensure fair comparison across methods producing clusterings at different levels of granularity, we additionally employed Element-Centric Similarity (ECS), a resolution-agnostic metric specifically designed to compare clusterings with differing numbers of clusters. For deconvolution-based methods, including RCTD, Cell2location, CARD, SpatialDWLS, SPOTlight, DestVI, and TACCO, the predicted cell type abundance matrix was converted to a discrete annotation by assigning each cell or spatial location to the cell type with the maximum estimated proportion. The resulting labels were used for all downstream performance evaluations.

### Cell type annotation methods Dimensionality-reduction-based framework

***CARD*** [9] is a deconvolution method which uses non-negative matrix factorization (NMF) to combine the gene-by-cell type data from a reference scRNAseq dataset with gene-by-spot spatial data. It also incorporates an autoregressive weighting step that favors similar cell type compositions among neighboring ST spots, mimicking spatial continuity in tissue. This extra step allows CARD to deconvolve ST data of different resolutions, while providing some robustness against mislabeled reference data.

### Optimal transport framework

***TACCO*** [11] is a modular and flexible framework for deconvolution or cell type annotation. By default, it applies an optimal transport (OT) method based on cell type probabilities to combine unlabeled spatial transcriptomic data with reference sequencing data. TACCO can also be easily configured to compare against a ground truth or exchange OT for non-negative least squares (NNLS).

### Regression-based framework

***SpatialDWLS*** [7] uses a reference scRNA-seq dataset to estimate cell type proportions in spatial transcriptomic spots. It assumes cell types can be identified by a characteristic marker gene and then applies dampened weighted least squares (DWLS) to minimize the difference between the model’s predicted and observed ST data. The dampening step allows the model to manipulate data with high variability in gene expression between spots.

***SPOTlight*** [8] utilizes non-negative matrix factorization (NMF) and non-negative least squares regression (NNLS) to deconvolute spatial transcriptomics data. The model identifies topics (latent representations of gene expression patterns) and computes a topic profile distribution for each spatial location. Each cell is then assigned a type that best matches its topic distribution. A notable advantage of SPOTlight is its ability to produce accurate results with small or shallowly sequenced single-cell RNA-seq datasets.

### Poisson distribution framework

***RCTD*** [32] fits a Poisson statistical model to describe each pixel as a linear combination of cell types, inferring cell type proportions through maximumlikelihood estimation (MLE). The resulting output is a matrix containing estimated cell type proportions for every spatial location. RCTD aims to improve on other supervised learning methods by accounting for the differences in gene capture rates arising from platform effects.

### Optimization-based framework

***Tangram*** [13] utilizes gradient-based optimization to map single-cell RNA-seq data of a tissue to spatial transcriptomic data of that same tissue. It combines cell density, gene expression by spot and total gene expression to form a detailed cost function and outputs and an estimate of cell type proportion by spot. By integrating histological and anatomical data into its analysis, Tangram can improve the resolution and quality of the input ST data.

### Anchor-based nearest neighbor framework

***Seurat*** [10] uses an anchor-based label transfer framework, first finding mutual nearest neighbors among a single-cell RNA-seq reference dataset and the query dataset to identify corresponding (“anchor”) cells. Each anchor pair is assigned a weight proportional to the strength of the match. Correction vectors are computed for each cell in the target dataset and used to refine the anchor weights, which are then passed to a weighted-vote classifier that outputs a cell type probability distribution and assigns the final label to each cell.

### Correlation-based framework

***SingleR*** [12] is a reference-based annotation method that assigns cell type labels to query cells by comparing their expression profiles with a labeled reference. It first identifies marker genes from pairwise comparisons between reference labels and computes Spearman correlation-based scores between each query cell and reference labels. The highest-scoring label is assigned to each cell, with an optional fine-tuning step that recalculates scores among closely related candidate labels using label-discriminative marker genes.

### Negative binomial distribution framework

***DestVI*** [31] is a Bayesian model based on negative binomial regression consisting of two components: scLVM (single-cell latent-variable model) and stLVM (spatial transcriptomic latent-variable model). scLVM learns latent representations from single-cell RNA-seq data using a conditional generative framework, and stLVM uses these representations to deconvolve spatial transcriptomics data. The resulting output is a matrix containing the predicted quantities of each cell type for every spatial location. An advantage of DestVI is its ability to process large datasets containing up to a million cells.

***Cell2location*** [1] is a Bayesian model that outputs a matrix containing the predicted count of each cell type for every spatial location. It uses a negative binomial regression model to compute a matrix of reference cell type signatures derived from single-cell RNA-seq data. These signatures are then used to decompose the spatial gene expression matrix into cell type specific contributions, outputting the estimated cell type abundances for each location.

### Deep learning-based framework

***Spatial-ID*** [29] is a deep learning-based reference-mapping framework for spatial cell type annotation. It uses annotated scRNA-seq data for model pretraining, and then incorporates both gene expression and spatial neighborhood structure to learn spatially informed latent embeddings. Based on these embeddings, Spatial-ID predicts cell type probabilities and assigns annotation labels to individual spatial locations.

***scANVI*** [14] is a semi-supervised deep generative model that utilizes variational inference. It is designed to infer remaining cell types in datasets where only a subset of cells has pre-existing labels, but can also be used for label transfer when given a singlecell RNA-seq reference dataset. scANVI first generates latent space embeddings for all cells, and then uses a Bayesian approach to annotate unlabeled cells based on their latent representations. The resulting output is a cell type probability distribution and a final predicted label for each unlabeled cell.

***GraphST*** [30] is a graph self-supervised contrastive learning framework for spatial clustering, multisample integration, and cell type deconvolution of spatial transcriptomics data. It combines graph neural networks with contrastive learning to model gene expression, spatial location, and local neighborhood context, encouraging spatially adjacent spots to have similar latent representations. For reference-based analysis, GraphST can transfer single-cell RNA-seq information onto spatial transcriptomics data to infer cell type composition or map reference cells to spatial locations.

### Foundation model

***scGPT*** [15] is a transcriptome foundation model pretrained on large-scale scRNA-seq data for diverse single-cell biology tasks. It learns generalizable representations of gene-expression programs and cell states that can be adapted to downstream cell type annotation. For label transfer, we considered one zeroshot mapping strategy and two supervised adaptation strategies for the pretrained scGPT model: reference mapping, head tuning, and partial fine-tuning. In the head-tuning setting, the pretrained transcriptomic encoder was kept fixed and only the cell type classification head was optimized using labeled reference cells, allowing the model to learn a dataset-specific label mapping while preserving the pretrained gene–cell representations. In the partial fine tuning setting, we initialized scGPT from the pretrained checkpoint and froze the non-transformer encoder components, including the gene/token embedding and related input encoders, while allowing the transformer encoder and the cell type classification head to be updated using labeled reference cells. This strategy retains key pretrained input representations while permitting the transformer backbone to adapt to the target reference dataset. Gene-expression profiles were normalized, discretized into expression bins, tokenized by gene identity, and passed through the model, where the CLS cell embedding was used for supervised cell type prediction. The trained model was then applied to query cells to generate cell type probability distributions and final predicted cell type labels. For scGPT reference mapping, reference and query cells were first embedded using the pretrained scGPT model without task specific fine tuning. Query cells were then mapped to the reference embedding space, and cell type labels were transferred from the nearest reference cells using a similarity-based nearest-neighbor voting strategy.

***scCello*** [27] is a transcriptome foundation model that incorporates hierarchical relationships between cell types through cell-ontology graphs. Its pretraining framework integrates gene-level expression modeling, intra-cellular objectives that encourage consistent representations among cells of the same type, and inter-cellular objectives that preserve ontological relationships across cell types. For label transfer, we adapted the pretrained scCello model using two supervised training strategies: head tuning and partial fine tuning. In the head tuning setting, the pretrained BERT-based transcriptomic backbone was frozen and only the task-specific cell type classification head was optimized using labeled reference cells. In the partial fine-tuning setting, we initialized scCello from the pretrained checkpoint, froze most pretrained backbone parameters, and updated only the final two encoder layers, the pooler, the embedding layer normalization parameters, and the classification head. The adapted model was trained with a supervised cell type classification objective on labeled reference cells and then applied to query cells to generate cell type probability distributions and final predicted labels.

***Nicheformer*** [16] is a transformer-based foundation model pretrained on both dissociated single-cell and spatial transcriptomics data to learn cell representations that capture spatial context. The pretraining corpus includes both human and mouse datasets, enabling the model to capture cross-species biological patterns. Cells are represented as gene-expression tokens ordered by normalized expression levels after adjustment for technology-dependent biases. For label transfer, we adapted the pretrained Nicheformer model using two supervised training strategies: head tuning and partial fine tuning. In the head-tuning setting, the pretrained backbone was frozen and only the task-specific prediction head was trained on labeled reference cells. In the partial fine-tuning setting, we initialized the model from the pretrained checkpoint and froze most pretrained parameters, including the embedding layers, positional embeddings, and the first nine encoder layers. The final three encoder layers and the prediction head were updated using labeled reference cells, with a lower learning rate for the unfrozen backbone layers than for the prediction head. The trained model was then applied to query cells to obtain class logits, which were converted to final predicted cell type labels by selecting the class with the highest score.

***SToFM*** [28] is a multi-scale spatial transcriptomics foundation model designed to learn cell representations by jointly integrating tissue-level morphology, local cellular microenvironment, and gene-expression information. It constructs sub-slices from each ST section to capture macro-, micro-, and gene-scale features, and then applies an SE(2) Transformer to generate spatially aware cell embeddings for downstream tasks such as tissue region segmentation and cell type annotation. We performed partial fine tuning by freezing most of the pretrained SToFM backbone while unfreezing the final three SE(2) Transformer layers and training them jointly with an MLP classification head.

### Unsupervised methods

***STdeconvolve*** [33] is an unsupervised method for deconvolving spatial transcriptomic data using a Latent Dirichlet Allocation (LDA) model. It treats each spatial transcriptomics capture pixel as a mixture of cell types, analogous to a document containing multiple topics in LDA. Through this approach, STdeconvolve simultaneously infers the transcriptional profile of each cell type and annotates each cell using the dominant component (hard label).

***SpaGCN*** [3] uses spatial transcriptomic and optional histological data to identify spatially variable genes and the patterns in which they vary. This unsupervised model uses graph convolution weighted by Euclidean distance to group sampled spots into spatial domains. Then it searches for a gene or combination of genes whose expression matches each spatial domain. This allows SpaGCN to find spatially variable genes and their gradients, which may lead to information about cell type gradients or tissue boundaries in the sample.

***BANKSY*** [34] constructs augmented feature vectors by concatenating each cell’s expression profile with neighborhood-averaged features and spatial feature gradients, thereby encoding high-order spatial dependencies. Clustering in this enriched feature space enables a unified framework for both finegrained cell typing and tissue-domain segmentation with high scalability.

### Evaluation metrics

#### Macro F1

To assess performance under class imbalance, we report the macro-averaged F1 score, which assigns equal weight to each class by averaging class-wise F1 scores [38]. This is the standard implementation used in common libraries such as scikitlearn. Although alternative definitions exist in the literature (e.g., averaging precision and recall before computing F1), they coincide only under certain conditions and are not used here. For each cell type *c*, we compute

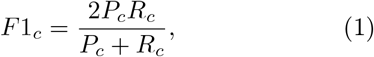

where *P*_*c*_ and *R*_*c*_ denote the precision and recall for class *c*, respectively. The overall macro-averaged F1 score is then defined as

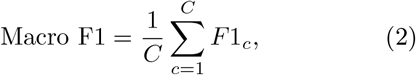

where *C* is the total number of cell groups. Compared with accuracy or micro-averaged metrics, F1-macro is more sensitive to rare classes and better reflects per-class annotation quality in datasets with imbalanced cell type compositions. The macro F1 score ranges from 0 to 1, with higher values indicating stronger agreement between predicted and ground-truth labels across classes.

#### Micro F1

To complement macro-averaged evaluation, we also report the micro-averaged F1 score, which pools true positives, false positives, and false negatives across classes before computing the F1 score [39, 38, 40]. Let TP_*c*_, FP_*c*_, and FN_*c*_ denote the counts for class *c*. We define

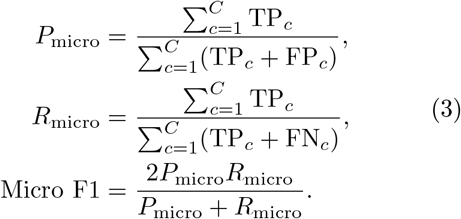

The micro F1 score ranges from 0 to 1, with higher values indicating better overall classification performance. In the single-label multi-class setting, micro F1 is equivalent to overall accuracy because it aggregates predictions across all classes before computing the F1 score.

#### Weighted F1

We additionally report the weighted F1 score, which averages class-wise F1 scores while weighting each class by its cell number [39, 41]. Specifically, let *n*_*c*_ denote the number of cells in class *c* and 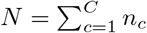. The weighted F1 score is defined as

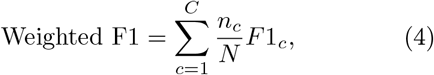

where 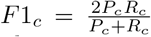 is the per-class F1 score. The weighted F1 score ranges from 0 to 1, with higher values indicating better classification performance. Compared to macro F1, weighted F1 down-weights rare cell types and is therefore less sensitive to class imbalance, while still reflecting per-class performance more explicitly than micro-averaged metrics [38].

#### Element-Centric Similarity (ECS)

To complement global agreement metrics (e.g., ARI/NMI) with a cell-level measure of label consistency, we quantified the similarity between the ground-truth labeling *a* and the method-assigned labeling *b* using element-centric similarity (ECS). Using the affinity-matrix formulation, for a labeling *a* over *N* cells, we construct an *N* × *N* affinity matrix *P*^*a*^, where 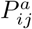 denotes the connection strength between cells *i* and *j*:

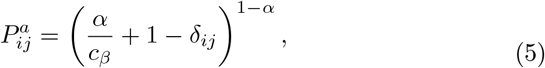

and

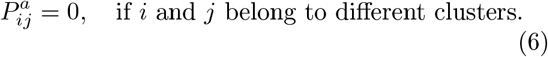

Here, *c*_*β*_ denotes the size of cluster *β, δ*_*ij*_ is the Kronecker delta, and *α* controls intra-cluster weighting (we used *α* = 0.9). We analogously construct *P*^*b*^ for labeling *b* [42].

ECS is then defined at the per-cell level by comparing the affinity profiles of each cell *k* between *P*^*a*^ and *P*^*b*^:

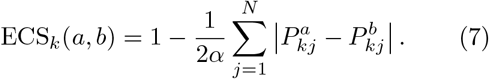

Finally, the overall pairwise similarity between labelings *a* and *b* is computed by averaging over cells:

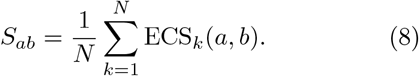

In our benchmarking, we report *S*_*ab*_ as a complementary metric to accuracy-based evaluations, as it captures cell-wise membership consistency and is particularly informative under cluster-size imbalance or when fine-grained subtypes exhibit transcriptional overlap. The resulting similarity score *S*_*ab*_ ranges from 0 to 1, where larger values indicate stronger celllevel membership consistency between the groundtruth and method-assigned labelings, and *S*_*ab*_ = 1 corresponds to identical label assignments.

#### Adjusted Rand Index (ARI)

Let *U* and *V* denote the predicted clustering and reference labels, respectively. The adjusted Rand index (ARI) [40] measures pairwise agreement between two partitions, corrected for chance, and is defined as

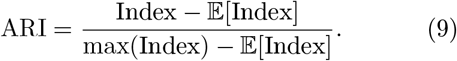

Here, Index denotes the Rand index, defined as the proportion of cell pairs that are either assigned to the same cluster in both *U* and *V* or assigned to different clusters in both *U* and *V*. The ARI ranges from 1 to 1, where − 1 indicates perfect agreement and 0 corresponds to the expected agreement under random labeling.

#### Normalized Mutual Information (NMI)

The normalized mutual information (NMI) [40] quantifies the shared information between the predicted clustering *U* and reference labels *V*, and is defined as

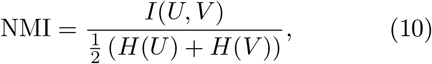

where *I*(*U, V*) denotes the mutual information and *H*(·) denotes entropy. The NMI ranges from 0 to 1, with higher values indicating stronger agreement.

#### Adjusted Mutual Information (AMI)

The adjusted mutual information (AMI) [40] corrects mutual information for chance agreement under a random labeling model and is defined as

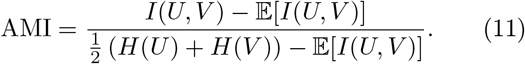

The AMI equals 1 for identical partitions and is approximately 0 under random labeling.

#### Average Silhouette Width (ASW)

The average silhouette width (ASW) [43] evaluates cluster compactness and separation based on pairwise distances between cell embeddings and is defined as

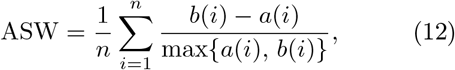

where *a*(*i*) is the average distance between cell *i* and all other cells in the same cluster, and *b*(*i*) is the minimum average distance to cells in other clusters. The ASW ranges from 1 to − 1, with higher values indicating better-defined clusters.

#### Average Biological Conservation Score (AvgBIO)

We use AvgBIO [15] as a summary metric of label-transfer quality. Higher values indicate better conservation of biologically meaningful structure, while low values suggest potential biological distortion. Given *K* label-consistency metrics 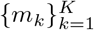 (all oriented so that larger is better, and scaled to [0, 1]), we define

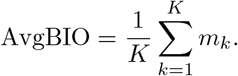

In our benchmark, we used

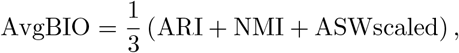

ASW was originally computed on the [-1, 1] scale and then linearly transformed to [0, 1] using (ASW+1)/2 before being included in AvgBIO. Thus, AvgBIO was defined as the average of NMI, ARI, and the rescaled ASW. AvgBIO ranges from 0 to 1, with higher values indicating better overall concordance between transferred labels and true cell identities. By integrating multiple complementary metrics, AvgBIO reduces over-reliance on any single metric.

#### Cohen’s d (score separability)

We used Cohen’s *d* [44] as a standardized effect-size metric to quantify how well each method’s binary cell type annotation separates a target cell type from non-targets in terms of marker-based enrichment scores. For each method and each cell type, we first computed a per-cell enrichment score *x* using the ground-truth marker gene set for that cell type. We then defined a binary prediction label *y* ∈ { 0, 1} from the method’s annotation, where *y* = 1 indicates the method annotated the cell as the target type and *y* = 0 otherwise. Cohen’s *d* was computed by comparing the enrichment-score distributions of the predicted-positive and predictednegative groups:

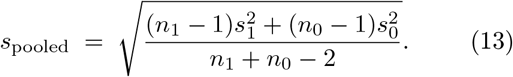

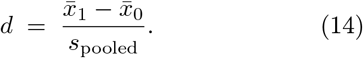

where 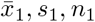 denote the mean, standard deviation (computed with Bessel’s correction, i.e., using denominator *n* − 1), and sample size of enrichment scores among cells with *y* = 1, and 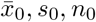 denote the corresponding quantities among cells with *y* = 0. Here, *s*_pooled_ denotes the pooled within-group standard deviation of enrichment scores across the predicted-positive and predicted-negative groups. Larger positive values of *d* indicate that cells annotated as the target type by the method have higher marker enrichment than cells annotated as non-targets, implying better score-based separation; values near zero indicate substantial overlap. We reported *d* as missing when either group had fewer than two cells or when either group’s sample variance was zero.

#### AUC-ROC (Area Under the Receiver Operating Characteristic curve)

For each method and each cell type *ct*, we computed a per-cell enrichment score *x* using the ground-truth marker gene set for *ct*, defined a binary prediction label *y* ∈ { 0, 1} where *y*_*i*_ = 1 if the method predicted cell *i* as *ct* (and 0 otherwise), and used the AUC-ROC which ranges from 0 to 1 to quantify how well the method’s predicted *ct* cells are ranked above non-*ct* cells by the marker-based enrichment score [45]:

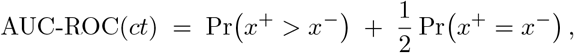

where *x*^+^ and *x*^−^ are enrichment scores from predicted-positive and predicted-negative cells, respectively.

### Enrichment analysis

We used single-sample gene set enrichment analysis (ssGSEA) [46], implemented in the GSVA R package [47], to score each cell against predefined marker gene sets derived from the ground-truth cell type annotations. For each annotation method and target cell type, we quantified concordance between predicted labels and the corresponding ground-truth marker score by calculating the area under the receiver operating characteristic curve (AUC-ROC) between the ssGSEA score vector and a binary indicator of whether each cell was assigned to that cell type. To assess pathway-level biological coherence, we next computed ssGSEA scores using the Reactome gene sets from MSigDB v2026.1.Hs [48, 49, 50]. For each ground-truth cell type, we selected the 10 Reactome pathways with the highest ssGSEA scores in the ground-truth population, and compared their enrichment across the corresponding method-predicted populations.

### Real Datasets

#### MERFISH mouse spinal cord

The mouse spinal cord MERFISH dataset comprises 18 spatial sections from five adult mice [24]. Each measurement unit is a cell. The dataset comprised 18 sections spanning the cervical, thoracic, lumbar, and sacral regions of the spinal cord, including four cervical, five thoracic, three lumbar, and six sacral sections. Cell types were jointly annotated based on literature-informed marker genes.

#### A harmonized atlas of single-nucleus RNA-seq mouse spinal cord

We used the harmonized postnatal mouse spinal cord cell type atlas curated by Russ et al [35]. as an external reference for annotating query MERFISH mouse spinal cord data. This atlas was constructed by preprocessing and integrating raw data from six publicly available postnatal mouse spinal cord single nucleus RNA-seq studies. The resulting harmonized reference comprises 52,623 nuclei with 23,901 quantified genes and provides curated annotations for 15 non-neuronal and 69 neuronal cell types [35]. Because cell type labeling framework differs between our MERFISH query dataset and the external single-nucleus RNA-seq atlas, we harmonized labels prior to evaluation. Specifically, we constructed a curated mapping dictionary to translate MERFISH ground-truth annotations into the corresponding atlas cell type names, enabling consistent label spaces for downstream metric computation. Cell types present in the MERFISH annotations but absent from the atlas (i.e., without a mapped counterpart) were excluded from quantitative evaluation; accordingly, all performance metrics were computed on the subset of cells whose ground-truth labels could be matched to the reference taxonomy. In cases where multiple MERFISH subtypes corresponded to a single atlas category, we performed many-to-one collapsing to the atlas label for evaluation.

#### Open-ST human metastatic lymph node

We included a human metastatic lymph node spatial transcriptomics dataset generated using the OpenST platform, which profiles tissues at single-cell resolution using a sequencing-based spatial transcriptomics workflow with whole transcriptome coverage. Cell type annotations provided in the original study were derived from canonical marker gene expression and further validated through histopathological assessment and cross-platform comparison with an imaging-based spatial transcriptomics platform (Xenium). The dataset comprises 15 annotated cell types, capturing diverse immune, stromal, and tumor-associated populations [25]. The human metastatic lymph node dataset consists of 19 serial tissue sections, spanning approximately 350 µm of tissue depth. For benchmarking, we selected section ID 6 as the reference dataset and section ID 19 as the testing dataset, balancing computational feasibility with biologically meaningful spatial and cellular complexity.

#### Six stages Xenium mouse renal injury and repair

We included a recently published multimodal mouse kidney dataset that combines high-resolution Xenium spatial transcriptomics with matched singlenucleus RNA-seq across six time points spanning renal ischemia–reperfusion injury (IRI) and subsequent repair [37]. This dataset profiled kidney tissues at six time points spanning injury and repair (Sham, Hour 4, Hour 12, Day 2, Day 14, and Week 6 post-injury), enabling systematic evaluation of cell type annotation performance under dynamic pathological conditions. Xenium in situ profiling was performed using a custom 300-gene panel, targeting canonical kidney cell type markers, injury-associated genes, and signaling molecules relevant to fibrosis and immune responses [2]. The Xenium dataset comprised over 1.3 million spatially resolved single cells, with cell identities annotated into 20 major kidney cell populations based on canonical marker genes. A matched single-nucleus RNA-seq dataset [37] spanning the same six disease stages was used as the reference for reference-based label transfer methods.

#### Stereo-seq developmental axolotl

We benchmarked methods on an axolotl telencephalon singlecell spatial transcriptomics dataset generated using Stereo-seq, a platform that provides spatially resolved transcriptomics at near single-cell resolution by assigning transcripts to segmented individual cells. The dataset spans both normal development and injury-induced regeneration, including six developmental stages and seven post-injury time points after lateral pallium resection [26]. Across these samples, the original study identified 33 developmental cell types and 28 regenerative cell types, capturing major neural and glial populations as well as transitional and regeneration-associated states. For our analysis, we selected stage 44, stage 54, and 5 days post-injury, which together represent early developmental neurogenesis, a more advanced developmental state with emerging spatial restriction of cell populations, and an early regenerative response after injury.

## Computation platform

All experiments were conducted on a computer server equipped with Intel Xeon W-2195 CPUs, running at 2.3 GHz, featuring 25MB of L3 cache and 36 CPU cores. The server was configured with 256GB of DDR4 memory operating at 2,666MHz.

For GPU configurations, the same server was used, equipped with four Quadro RTX A6000 cards, each providing 48GB of memory and 4608 CUDA cores.

## Authors’ contributions

X.M.Z. and S.M. conceived and led this work. Y.Z. and X.M.Z. designed the framework. Y.Z., Y.H., M.B.X., and H.Q. performed analyses. Y.Z., S.M., and X.M.Z. wrote the manuscript. All authors reviewed the manuscript.

## Data availability

All code, tutorials, evaluation scripts, and related data files are freely available on GitHub https://github.com/maiziezhoulab/Benchmark_ST_cell_type_annotation.git and on Zenodo with DOI: https://doi.org/10.5281/zenodo.20720105 under the MIT license. A summary of the data is shown in Table 2. Dataset 1 consists of 18 mouse spinal cord sections, available at https://doi.org/10.5281/zenodo.18039571 with annotation [24]. Dataset 2 includes mouse spinal cord single-nucleus RNA-seq data available at https://www.ncbi.nlm.nih.gov/geo/query/acc.cgi?acc=GSE158380 with annotation [35]. Dataset 3 contains 2 human metastatic lymph node at https://www.ncbi.nlm.nih.gov/geo/query/acc.cgi with annotation [25]. Dataset 4 includes twelve slices of male mouse kidneys across six stages of renal injury and repair available at https://www.ncbi.nlm.nih.gov/geo/query/acc.cgi?acc=GSE269719 with annotation [2]. Dataset 5 is the mouse single-nucleus RNA-seq of acute kidney injury available at https://www.ncbi.nlm.nih.gov/geo/query/acc.cgi?acc=GSE139107 with annotation [37]. Dataset 6 is a Stereo-seq spatial transcriptomics dataset of axolotl telencephalon development and regeneration available at https://db.cngb.org/stomics/artista with annotation [26].

## Ethics approval and consent to participate

Not applicable.

## Consent for publication

Not applicable.

