## Supplemental Figures for "Benchmarking cell type annotation in spatial transcriptomics: resolving cellular hierarchies, biological fidelity, and dynamic cell states"

Zhu et al.

### 1 Supplementary figures

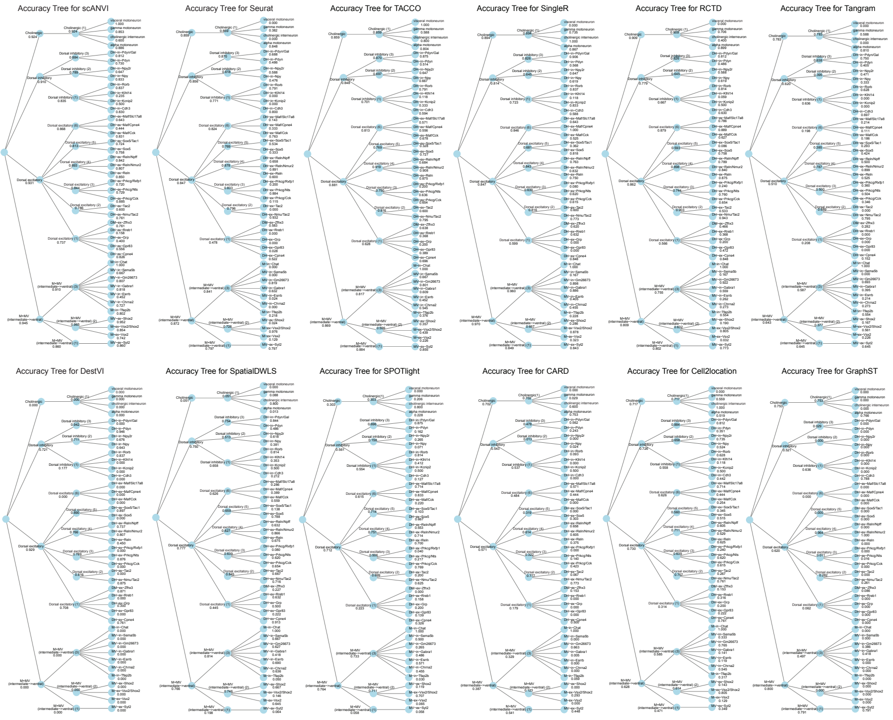

Figure S1: **Neuronal cell type hierarchy and classification accuracy produced different methods.** Cell identities are organized hierarchically from broad neuronal classes to increasingly fine-grained subtypes. Internal nodes represent intermediate cell type categories, whereas leaf nodes represent terminal cell types. Values shown beside each node indicate the classification accuracy by each method at that level of the hierarchy. This tree-based visualization assesses whether each method captures only coarse lineage structure or also preserves fine-grained subtype specificity.

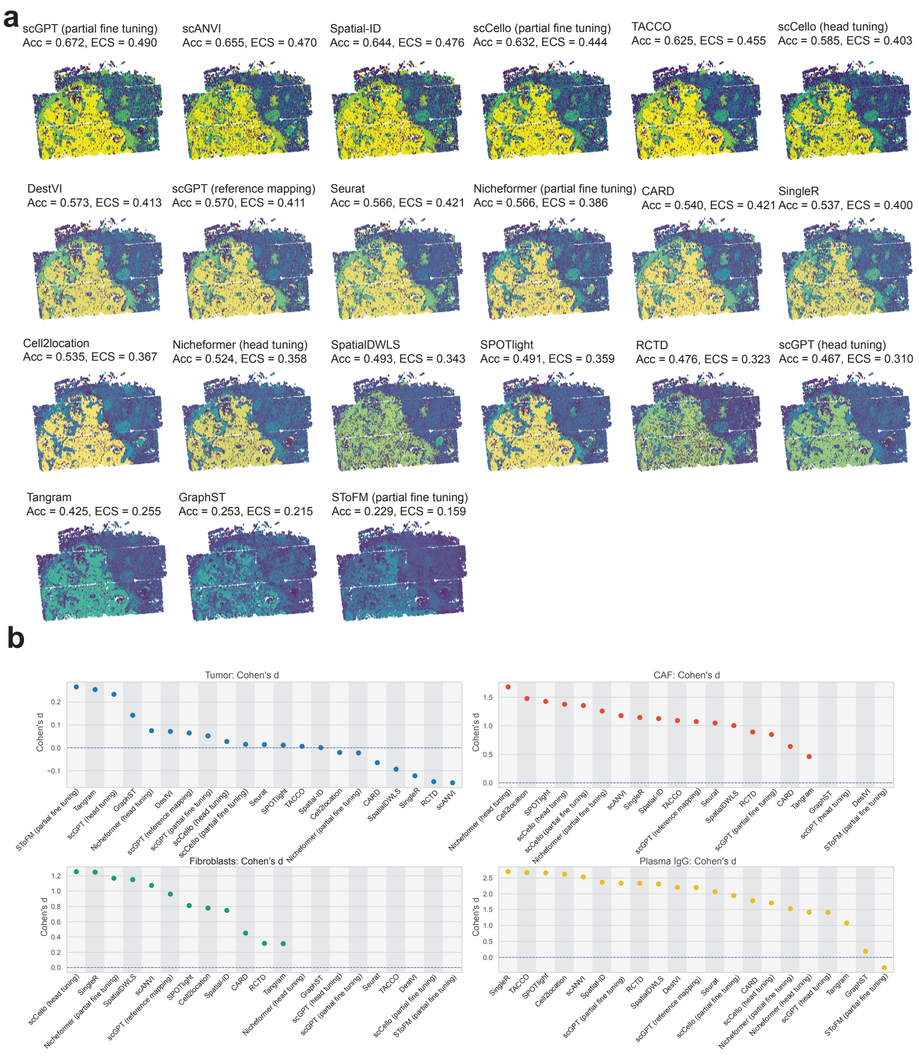

Figure S2: **Spatial label consistency and marker-program separability across annotation methods (a)** Spatial maps of predicted cell type labels for each annotation method. Accuracy and element-centric similarity (ECS) are shown above each map. ECS was computed by comparing per-cell affinity profiles between ground-truth and predicted labelings, providing a cell-level measure of label membership consistency. **(b)** Cohen's d analysis of marker-score separability for Tumor cells, CAF, Fibroblasts, and Plasma IgG cells. For each method, enrichment scores of the target marker program were compared between cells predicted as the target cell type and predicted non-target cells. Larger positive values indicate stronger marker enrichment among predicted-positive cells, whereas values near zero or below zero indicate weak or inconsistent biological separation.

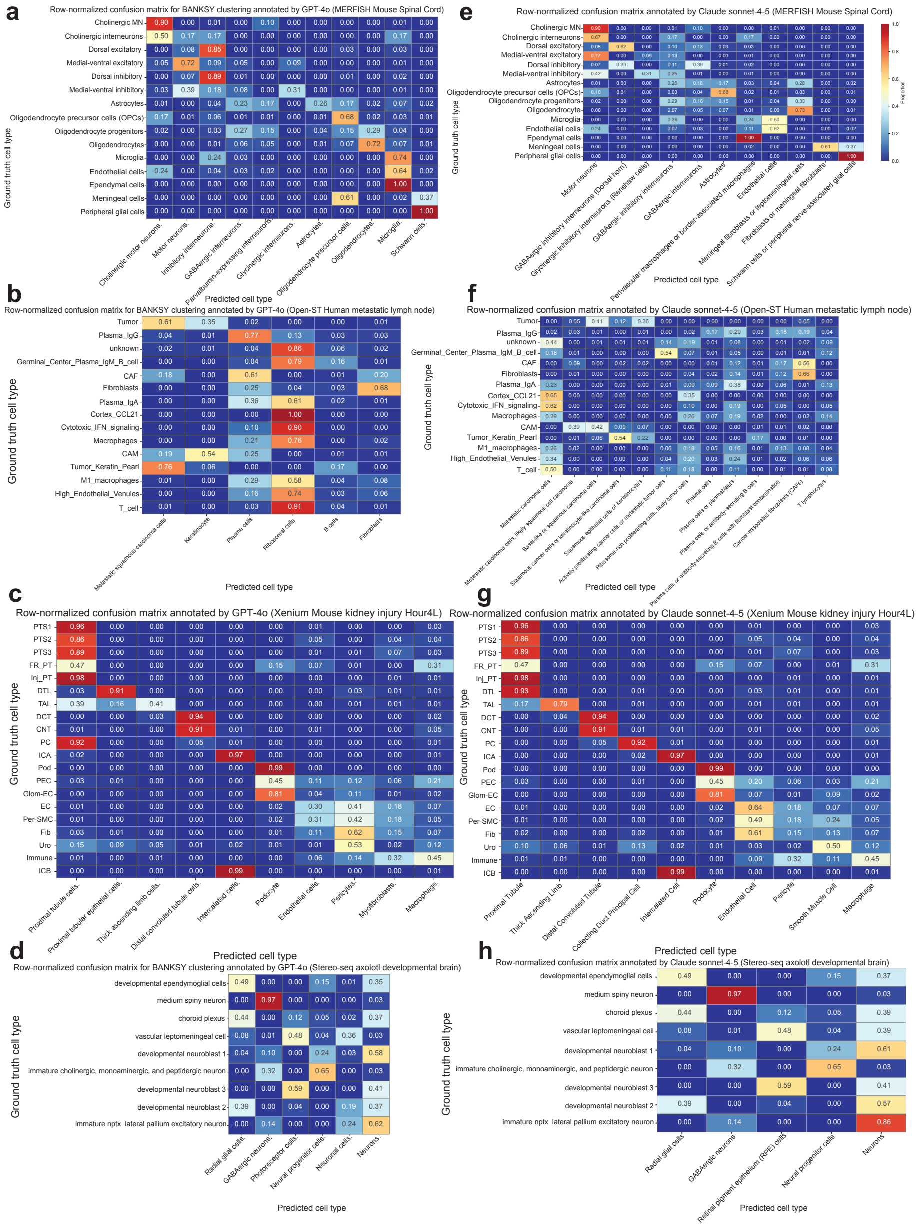

Figure S3: **Comparison of cell type annotation methods using LLM-based annotation. (a–h)** Row-normalized confusion matrices for LLM-based cell type annotation across four spatial transcriptomics datasets. For each dataset, de novo clustering was first performed using BANKSY, followed by cluster-level annotation with GPT-4o or Claude Sonnet 4-5. Both ground-truth and predicted cell type labels were ordered by biological lineage.
